# Absence seizures and sleep-wake abnormalities in a rat model of *GRIN2B* neurodevelopmental disorder

**DOI:** 10.1101/2024.02.27.582289

**Authors:** Katerina Hristova, Melissa C. M. Fasol, Niamh McLaughlin, Sarfaraz Nawaz, Mehmet Taskiran, Ingrid Buller-Peralta, Alejandro Bassi, Adrian Ocampo-Garces, Javier Escudero, Peter C. Kind, Alfredo Gonzalez-Sulser

## Abstract

Pathogenic mutations in *GRIN2B* are an important cause of severe neurodevelopmental disorders resulting in epilepsy, autism and intellectual disability. *GRIN2B* encodes the GluN2B subunit of N-methyl-D-aspartate receptors (NMDARs), which are ionotropic glutamate receptors critical for normal development of the nervous system and synaptic plasticity.

Here, we characterized a novel *Grin2b* heterozygous knockout rat model with 24-hour EEG recordings. We found rats heterozygous for the deletion (*Grin2b^+/-^*) had a higher incidence of spontaneous spike and wave discharges, the electrographic correlate of absence seizures, than wild-type rats (*Grin2b^+/+^*). Spike and wave discharges were longer in duration and displayed higher overall spectral power in *Grin2b^+/-^* when compared to those in *Grin2b^+/+^* animals. Heterozygous mutant rats also had abnormal sleep-wake brain state dynamics over the circadian cycle. Specifically, we identified a reduction in total rapid eye movement sleep and, altered distributions of non-rapid eye movement sleep and wake epochs, when compared to controls. This was accompanied by an increase in overall spectral power during non-rapid eye movement sleep in *Grin2b^+/-^*. The sleep-wake phenotypes were largely uncorrelated to the incidence of spike and wave discharges.

We then tested the antiseizure efficacy of ethosuximide, a T-type voltage-gated calcium channel blocker used in the treatment of absence seizures, and memantine, a noncompetitive NMDAR antagonist currently explored as a mono or adjunctive treatment option in NMDAR related neurodevelopmental disorders. Ethosuximide reduced the number and duration of spike and wave discharges, while memantine did not affect the number of spike and wave discharges but reduced their duration.

These results highlight two potential therapeutic options for *GRIN2B* related epilepsy. Our data shows the new rat *Grin2b* haploinsufficiency model exhibits clinically relevant phenotypes. As such, it could prove crucial in deciphering underlying pathological mechanisms and developing new therapeutically translatable strategies for *GRIN2B* neurodevelopmental disorders.

## Introduction

*GRIN2B* pathogenic variants cause multiple neurodevelopmental clinical phenotypes such as epilepsy, autism spectrum disorder (ASD), intellectual disability, sleep impairments and movement abnormalities (Endele et al., 2010; Epi4K Consortium., 2013; Freunscht et al., 2013; O’Roak et al., 2011; Platzer et al., 2017). An estimated 52% of patients with *GRIN2B*-related neurodevelopmental disorder present with severe epilepsy, which is refractory to therapy in half of diagnosed cases (Platzer et al., 2017). A variety of seizure types occur in patients including infantile spasms and generalized tonic-clonic, focal, and absence seizures (Epi4K Consortium., 2013; Platzer et al., 2017). De novo *GRIN2B* variants are one of the primary monogenic causes of epileptic encephalopathies, in which epilepsy is thought to worsen neurodevelopmental comorbidities (Epi4K Consortium., 2013; Lemke et al., 2014; Platzer et al., 2017; Scheffer et al., 2017; Smigiel et al., 2016). Dysfunctional sleep is common amongst individuals with GRIN2B mutations, and parental reports indicate at least 60% of patients struggle initiating and maintaining sleep, have breathing irregularities during sleep and suffer with daytime somnolence (Freunscht et al., 2013; Platzer et al., 2017; Simons Searchlight, 2021). The predicted incidence of *GRIN2B* pathogenic mutations is 5.91 per 100.000 births (Lemke, 2020), making it the most prevalent disorder associated with the N-methyl-D-aspartate receptor (NMDAR) coding *GRIN* genes.

NMDARs are sodium, potassium and calcium permeable tetrameric ligand-gated ion channels, which play a critical role in glutamatergic transmission, synaptic plasticity and the development of the nervous system (Paoletti et al., 2013; Traynelis et al., 2010). NMDARs are composed of two obligate glycine binding GluN1 subunits and either two GluN2 subunits, which bind to glutamate, or two GluN3 subunits, which bind to glycine (Cavara et al., 2009; Chatterton et al., 2002). There are four paralogs for GluN2 (GluN2(A-D)), which display differential expression profiles across development and result in varied channel activity dynamics (Paoletti et al., 2013; Traynelis et al., 2010).

*GRIN2B* encodes GluN2B, which is highly expressed in prenatal and early postnatal periods (Paoletti et al., 2013; Traynelis et al., 2010). Later in development the expression of GluN2A, which is encoded by *GRIN2A*, is increased. GluN2B containing receptors have a lower open probability, slower deactivation kinetics and an increased sensitivity to glutamate and glycine compared to GluN2A containing receptors (Paoletti et al., 2013; Wyllie et al., 2013). In the context of neurodevelopmental disorders, rodent in vitro data demonstrates that pathogenic *GRIN2B* variants alter NMDAR function (Kellner et al., 2021), neuronal migration (H. Jiang et al., 2015), dendrite morphogenesis (Bahry et al., 2021; Sceniak et al., 2019) and synaptic density (Liu et al., 2017; Sceniak et al., 2019), which likely contribute to the aberrant circuit properties that manifest in *GRIN2B*-related neurodevelopmental disorders.

To test the role of GluN2B on brain network function and the sleep-wake cycle, we performed chronic *in vivo* wireless EEG recordings in a novel rat model of *Grin2b* haploinsufficiency. We identify spontaneously occurring absence seizures, abnormal sleep-wake brain state patterns, and altered spectral properties in mutant animals. We also test the effects of clinically relevant pharmacological treatments on absence seizures.

## Materials and methods

### Animals

All animal procedures were undertaken in accordance with the University of Edinburgh animal welfare committee regulations and were performed under a United Kingdom Home Office project license. Long Evans *Grin2b* heterozygous knockout rats were generated by the Medical College of Wisconsin gene Editing Rat Resource Center with support from the Simons Foundation Autism Research Initiative (LE-Grin2bem1Mcwi, RRID:RGD_14394515), and *Grin2b* heterozygous knockouts are hereafter referred to as *Grin2b^+/-^*, while wild-type littermates are *Grin2b^+/+^*. Pups were bred in-house and weaned from their dams at postnatal day 22. They were subsequently housed in mixed genotype cages with wild-type littermates (two to six rats per cage) with a 12-hour/12-hour light-dark cycle with ad libitum access to water and food. Following surgery, animals were single housed. Rats were genotyped by polymerase chain reaction.

### Synaptosome preparation

Adult 19 week-old *Grin2b^+/-^* (six male, two female) and *Grin2b^+/+^* (three male, five female) rats were anesthetised with isoflurane and decapitated. The hippocampus and somatosensory cortex from each hemisphere were quickly dissected in slushy PBS 1X, snap frozen and stored at −80 °C. The discontinuous Percoll-density gradient (23% bottom, 10% middle, 3% uppermost) was made prior to homogenisation on the preparation day. The tissue was quickly thawed at 37 °C and homogenized in ice-cold homogenisation buffer (1x sucrose/EDTA, pH7.4) using 5-6 up-and-down strokes of a pre-chilled Teflon glass with a motorized homogenizer (Dunkley et al., 2008) followed by centrifugation at 2800 rpm for 10 min at 4°C. The supernatant was added gently on 3% Percoll-sucrose (Percoll, P1644, Sigma Aldrich, UK) and centrifuged at 20,000 rpm for 8 min at 4°C. The fraction between 23% and 10% was collected and re-suspended in HEPES-Buffered-Krebs (HBK; in mM: 118.5 NaCl, 4.7 KCl, 1.18 MgSO_4_, 10 Glucose, 1 Na2HPO_4_, 20 HEPES, pH 7.4 balanced with Trizma) followed by centrifugation at 13,000 rpm for 15 min at 4°C. The pellet (pure synaptosomes) was dissolved in RIPA buffer with protease inhibitor cocktail (5056489001, Roche, Switzerland) and phosphatase inhibitor cocktails II and III, (P5726 and P0044 respectively, Sigma Aldrich, United Kingdom). Protein concentration was determined with the MicroBCA Assay kit (Pierce BCA protein estimation kit, 23225, ThermoFisher Scientific, United Kingdom).

### Western Blot

Approximately 15 μg of synaptosome protein was separated on a precast gradient gel (4-15% Mini-PROTEAN® TGX™ Precast Protein Gels, 4561086, BioRad, United Kingdom) and transferred to nitrocellulose membrane (Amersham™ Protran® Western Blotting Membrane, Nitrocellulose, GE10600002, Sigma Aldrich, United Kingdom) using a Bio-Rad transfer apparatus. Total proteins were stained with a reversible protein stain kit (Memcode 24580, Thermo Fisher Scientific, United Kingdom) according to the manufacturer’s instructions. After removing the stain, membranes were blocked with 1:1 TBS1X: Odyssey Blocking Buffer (P/N-927-50003, LI-COR Biotech.) for an hour at room temperature, followed by overnight incubation with primary antibodies (NMDAR2A-1: 1000, #ab169873, Abcam; NMDAR2B-1:1000, #610417, BD Biosciences; PSD95-1:2000, #76115, Abcam; GluR1-1:1000, #MAB2263, Millipore) at 4°C. Membranes were washed with TBST1X (0.1% Tween 20), and incubated for an hour at room temperature with secondary antibodies (IRDye 680RD Donkey anti-rabbit IgG-1:5,000, #P/N 926-68073; IRDye 800CW Donkey anti-mouse IgG - 1:5,000, #P/N 926-32212, LI-COR Biotechnology, United Kingdom). Membranes were washed with TBST1X, dried and digitally scanned using an Odyssey M Imager, LI-COR, UK Ltd. Odyssey software, Licor Image Studio Lite (LICOR Biosciences, United Kingdom) was used to quantify individual bands. Data was normalised to respective total protein and normalised to wild-type levels.

### Surgery

Adult rats were anaesthetised with isoflurane and mounted on a stereotaxic frame (David Kopf Instruments, United States). Animals received subcutaneous Rimadyl Small Animal Solution (5 mg/kg, Zoetis, United states) for analgesia and 2.5 ml of sterile 0.9% saline for rehydration.

For surface EEG grid implantation, two holding screws for structural support (4 mm rostral, ± 0.5 mm lateral relative to bregma) and one ground screw over the cerebellum (−11.5 mm rostral, 0.5 mm lateral relative to bregma) (Yahata Neji, M1 Pan Head Stainless Steel Cross, RS Components, United Kingdom) were attached. 16-channel EEG surface grids with two integrated electromyogram (EMG) leads (Custom H16-Rat EEG16_Functional-NeuroNexus, United States) were placed over the skull with the EEG surface grid plus-symbol reference point aligned over bregma. Silver paint was used for connecting the ground screw with the grid. The implant was fixed to the skull using dental cement (Simplex Rapid, Kemdent, United Kingdom). A 21 G sterile needle tip was used to guide EMG wires (2.5 cm in length with 0.5 cm striped of insulation) into the neck muscle, after which the incision was sutured. Fifteen 16–9 week-old *Grin2b^+/-^*(eight male, seven female) and nine *Grin2b^+/+^* (three male, six female) animals were recorded and used for analyses. Four *Grin2b^+/-^*and nine *Grin2b^+/+^* rats that were descendants of one specific wild-type founder rat were excluded as they exhibited higher rates of absence seizures than descendants from other founders, which obfuscated genotype differences. Wild-type animals descended from the specific founder displayed between 1 and 1117 SWDs in 24 hours, on average 481 seizures over the rest of the population. Our current *Grin2b^+/+^* rats (decedents from multiple founders) do not display such a high prevalence of SWDs. Sample size estimates were not performed as these were the first recordings from this rat line.

For 2-channel EEG recordings for pharmacology experiments, bilateral screws were implanted to provide structural support (4 mm rostral, ± 0.5 mm lateral relative to bregma). Two ground screws were attached over the cerebellum (−11.5 mm rostral, ± 0.5 mm lateral relative to bregma), and two bilateral electrode screws were fixed over the primary somatosensory cortex (- 2.5 mm rostral, ± 4.5 lateral relative to bregma) (Yahata Neji, M1 Pan Head Stainless Steel Cross, RS Components). Electrode and ground screws were connected to an electronic interface board (EIB-16, Neuralynx) via silver wire soldered to each component (Teflon Coated Silver Wire, World Precision Instruments). The implant was fixed to the skull using dental cement (Simplex Rapid, Kemdent, United Kingdom). Twelve 21–26 week-old *Grin2b^+/-^* (ten male, two female) and five *Grin2b^+/+^* (one male, four female) animals were recorded and used in pharmacology experiments. A further four *Grin2b^+/+^*rats were implanted and recorded, but were excluded from pharmacological experiments as they did not display seizures during baseline recording days.

Site specific drug infusions were conducted specifically on *Grin2b^+/-^* rats. For drug infusion cannula implantation, two craniotomies were drilled bilaterally over the rostral reticular thalamic nucleus (nRT) (- 1.45 mm rostral, ± 2.2 lateral relative to bregma). The dura was removed from each craniotomy and a single 26 G sterile guide cannula (diameter = 0.46 mm, P1 Technologies) was slowly inserted to target the nRT (−5 mm ventral from the brain surface). Guide cannulas were cemented in place using UV activated cement (3M Relyx Unicem 2 Automix, Henry Schein) and dummy cannulas were placed into guides to ensure guides remained contaminant and blockage free. EEG and EMG components were then implanted as described above (2-Channel EEG). Four 20 week-old *Grin2b^+/-^* animals were implanted and used in site specific drug infusion experiments. An additional *Grin2b^+/-^* rat was implanted with cannulas, but was excluded from the final drug infusion analysis as the animal did not display SWDs during recording days.

### EEG recording

All animals were given a minimum of 1-week post-surgery recovery before recordings. Rats were habituated to the recording room for at least 24 hours. Animals were briefly anaesthetised with isoflurane to connect implants to wireless amplifiers, and subsequently recorded for 72 to 96 hours (for 24-hour sleep and seizure analysis) or for 4 hours (pharmacology experiments) using the TaiNi wireless multichannel recording system (Tainitec, United Kingdom) (Z. Jiang et al., 2017) at a sampling rate of 250.4Hz. Experimenters were blind to genotype.

### Absence seizure detection and sleep-wake scoring

The electrographic correlate of absence seizures, spike and wave discharges (SWDs), were characterized by periodic high-amplitude oscillations between 5 and 10 Hz (Meeren et al., 2002), which coincided with a spontaneous arrest of animal movement. We first utilized a previously published automatic SWD detector algorithm to quantify SWDs, which uses identification of harmonic peaks at 5-10 Hz in the power spectra to label seizure periods (Buller-Peralta et al., 2022; Katsanevaki et al., 2024). The code used for analysis is available at Zenodo and Github: https://zenodo.org/records/12700972.

As previously described, in summary, the SWD detection method consisted of the following: Spectral analysis revealed that visually scored SWDs behave as a high energy echo of a fundamental frequency (f0) located on the 5–10 Hz theta band, that resonates in several periodic harmonics across the frequency spectrum. This oscillating spectral structure of SWDs resembles a periodic waveform, allowing its automatic identification through a cepstral 1 analysis approach (Childers et al., 1977) by searching for a high amplitude peak located on a frequency band of interest. We applied an automated SWD seizure detection algorithm to voltage traces from the EEG grid electrode lead overlaid approximately on primary somatosensory cortex (right hemisphere, AP −3.0 mm and ML 2.8 mm from bregma), as, by visual assessment, was the channel most frequently associated with high amplitude SWDs across animals. After deconvolving the raw signal using a Fast Fourier Transform (number of tapers = 5), a logarithm was applied to obtain the magnitude. The signal could then be treated as semi-periodic so that the inverse Fast

Fourier Transform could be applied to obtain the cepstrum and reveal the period of the fundamental frequency (f0) as a spike in a pseudo-time domain frequency. After obtaining the cepstrum for the entire EEG recording (in sliding windows of 0.2 sec), peak power cepstrum values within the relevant frequency range (5–10 Hz) were identified and normalized by their absolute maximum. The resulting vector was transformed into z-scores to homogenise possible power differences between recordings that could distort seizure threshold identification. A threshold of ≥ 2.2 x 10^-5^ standard deviations was set by comparing the values of visually scored seizures against other high magnitude noise that resulted in false positives. 0.2 sec time windows were time-stamped as seizures when z-scored peak cepstral power in the theta band was greater than or equal to the established standard deviation threshold.

After SWD times were identified, a custom-made automated sleep scoring algorithm, based on a previously published method (Madrid-López et al., 2017), which clusters EEG and EMG recording epochs into corresponding sleep-wake states based on the spectral power at specific frequency bands associated with those states, was used to assign epochs to one of three brain states: wake, rapid eye movement (REM) sleep and non-rapid eye movement (NREM) sleep. Traces from EEG channels overlying primary somatosensory cortex on either brain hemisphere and the two EMG channels were visually inspected for artefacts. The EEG and EMG channels with fewer artefacts were chosen for analysis and if both EEG or EMG traces had similar noise levels, one channel for each modality was randomly chosen for analysis. Only one animal was excluded due to complete signal loss due to poor grounding. Spectral logarithmic EEG and EMG power was calculated for non-overlapping 5 second epochs in the 0.2–125 Hz frequency range using the *multitaper* package (Rahim et al., 2014) for R (R Core Team, 2022), using a non-overlapping window-size of 0.2 Hz half-time and bandwidth product of three. Each epoch was plotted in a three-dimensional metric space with values representing theta EEG power (maximum of the power in the 6.0 – 8.2 Hz range), delta and sigma power (1–20 Hz mean power, excluding the theta range), and neck muscle activity power (EMG power in the 60–90 Hz range). For clustering into brain states, centroids were estimated using an iterative algorithm that maximizes the local density by segregating the epochs based on the following characteristics: NREM sleep displayed high delta and sigma power with low EMG power, REM sleep displayed high theta power and low EMG power, and wake displayed high EMG power and low delta and sigma power. The distribution of clusters around the estimated centroid were approximated by Gaussian models assuming ellipsoidal shapes. The covariance matrix of the Gaussian models was used to calculate the Mahalanobis distance from a given epoch to each cluster. The epochs are assigned to a sleep stage according to a minimum Mahalanobis distance to the respective cluster.

All epochs identified to contain SWDs were classed separately from the other three brain states. An additional peak detection algorithm (van Brakel, 2014) was required to identify SWDs during wake epochs. This algorithm utilises z-scores to identify peaks by calculating a moving mean and flagging datapoints that deviate from the moving mean by a given threshold. If peaks were identified in both 5–9 Hz and 10–18 Hz, this was indicative of the presence of harmonics, which are evident in SWD epochs. After automated scoring of all additional SWD epochs the sleep quantification was adjusted accordingly. SWD distribution across wake and REM was determined by evaluating the first 5 second epoch prior to a seizure occurring, while the first 30 seconds (six epochs) prior were used to assess SWD initiation in NREM and wake-NREM transitions.

After initial brain state scoring, further processing was performed on the data to remove epochs with excessive (larger than 3000 mV) noise. A third-order bandpass Butterworth filter (0.2–100 Hz) was applied to the full-length raw data. 5 second epochs were extracted from the filtered recording and discarded if any data point exceeded 3000 mV, which was beyond the range of physiological activity.

To calculate the power spectral density, brain state epoch averages were Hanning-tapered and Fourier transformed using the SciPy welch method (with 50% overlapping windows). The spectral slope, a regression-based fitting measurement which describes the rate of change of the EEG power spectra, was calculated between 1–48 Hz using the NumPy polyfit function (Fasol et al., 2023). Epochs that were outside of the visually selected threshold of −5 log_10_(mV^2^)/Hz for the spectral slope and 500 mV for the spectral offset were removed from further analyses. Spectral power for each epoch was baseline-corrected by normalizing to the average spectral power across REM, NREM, and wake for each animal.

We validated automated sleep scoring performance by comparing the automatically generated vigilance state output with visual EEG state scoring of identical 5 second epochs. Researchers visually scoring were blind to automated output. We manually scored 4 hours of EEG in seventeen animals, between 10:00 – 14:00 o’clock, when approximately equal distributions of wake, NREM and REM epochs were observed. Percent agreement and Cohen’s kappa were calculated to assess interrater reliability between visual and automatic scoring of brain states.(McHugh, 2012) In our datasets, agreement between visual and automatic scoring was 88.1%, 87.5% and 95% for REM, NREM and wake states respectively. Overall agreement between the scoring methods was 90.8%. The kappa coefficient was 0.83 (± 0.002 SE) (Supplementary Fig. 1, Supplementary Table 1).

SWDs and brain state data was analysed over 48 hours starting at zeitgeber time 07:00 am of the day after connection, under a 12-hour/12-hour light–dark schedule. Values per hour over the two days were then averaged over the 24-hour period. The automated sleep scoring code is available at https://github.com/Gonzalez-Sulser-Team/AUTOMATIC-SLEEP-SCORER.

### Pharmacology experiments

For systemic pharmacology experiments, time-matched 2-hour periods from 11:00 to 13:00 o’clock across days, were used to analyse the effects of pharmacological treatments on SWDs. Treatment was administered 30 minutes prior to the start of the recorded and analysed period (at 10:30 am) and total recording durations for each experimental day did not exceed 4 hours. Drugs were dissolved in 1 ml of sterile 0.9% saline and rats received treatment based on their weight. Each animal received either intraperitoneal ethosuximide (100 mg/kg, Merck, Germany), memantine (5 mg/kg, Enzo Life Sciences, United States), or saline as a vehicle control on individual treatment days (Fig 6A), between baseline and washout days, with 24 hours between each experimental day (baseline, drug and washout) (Fig 6A). The half-lives of ethosuximide and memantine have been reported to be 10-16 hours and less than 4 hours respectively in rats (Beconi et al., 2011; Löscher, 2007), suggesting that a 24-hour washout is sufficient to record a new baseline period. Treatment order (including saline) was randomized and systematically weighted so that each possible permutation of treatment order occurred across an equal number of animals to minimize the risk of synergistic effects between drugs. SWD number for each treatment per individual rat is normalized to baseline SWDs and data is presented as a percentage of the animal’s baseline SWDs. These percentages are then compared statistically across the population. Experimenters were blinded to treatment.

In a separate experiment, we infused ethosuximide and memantine into the nRT, which is thought to be involved in SWD activity in other rodent models of absence seizures and patients (Crunelli et al., 2020; Lindquist et al., 2023). Drug preparation, administration and timeline followed was as described in systemic experiments above. A total volume of 1 μl was infused for each treatment, with 0.5 μl administered per hemisphere using an infusion cannula connected by polyethylene tubing to a 1μl Hamilton syringe. Injected volume (0.5 μl) and flow rate (10 nl/s) were controlled by a precision pump (World Precision Instruments, Hertfordshire, United Kingdom). During treatment administration rats were put under mild isoflurane anaesthesia, maintained between 1.5-2.5%, and the procedure was performed under a microscope to ensure infusion was effectively achieved. Drug infusions, under anaesthesia did not exceed 10 minutes (including brief anaesthetic induction). A 1-hour period between 11:00 to 12:00 o’clock across days, was used to analyse the effects of pharmacological treatments on SWDs. Based off the pharmacokinetic two-compartment model, the half-life for isoflurane elimination from the brain of rabbits is 26 minutes (Wyrwicz et al., 1987). Since the rat metabolism is slightly faster than that of rabbits (Subcommittee on Laboratory Animal Nutrition et al., 1995), and isoflurane exposure during infusions was short and at low concentrations, we reasoned that at 30 minutes most of the isoflurane is eliminated from the rat brain, while the following 1-hour period would still allow for observation of treatment effects. Treatment randomisation and SWD normalisation followed the same methods as for systemic administration and experimenters were blinded to treatment and genotype.

### Statistical analyses

Normality and homoscedasticity for all data were estimated by Shapiro–Wilk and Levene’s tests (rejection value set at < 0.05), and based on these, comparisons between genotypes were made using two-sample unpaired t-test, Welch’s two-sample t-test or Wilcoxon rank sum test. Two-way ANOVA tests were performed where more than two groups were compared (genotype and frequency). We use a Linear Mixed Model approach for time series analysis, where multiple measures were taken from the same subjects. Wherever appropriate, these were followed by *post hoc* Tukey’s multiple comparisons. Pearson correlation was used to test the relationship of SWDs with brain state abnormalities. Analyses were conducted in R (R Core Team, 2022), RStudio (Posit team, 2022), using the *ggplot2*, *car* and *lmerTest* packages (Fox & Weisberg, 2019; Kuznetsova et al., 2017; Wickham, 2016). All the data in the figures were presented as mean ± SEM, and *P* < 0.05 was considered as statistically significant and indicated with *\*P* < 0.05, *\*\*P* < 0.01, *\*\*\*P* < 0.001, *\*\*\*\*P <* 0.0001.

## Results

### Gene expression in a novel rat model of *GRIN2B* neurodevelopmental disorder

We utilized a novel model of *GRIN2B* related neurodevelopmental disorder generated by the Medical College of Wisconsin Gene Editing Rat Resource Centre, in which CRISPR/Cas9 was used to introduce a mutation in the *Grin2b* gene of Long-Evans rats. Protein expression was confirmed by immunoblotting somatosensory cortex (Fig. 1A and Supplementary Fig. 3A) and hippocampus (Supplementary Fig. 2A, Supplementary Fig. 3C) homogenates, and found to be located at synapses by immunoblotting of somatosensory and hippocampal synaptosome fractions (Fig. 1B and Supplementary Fig. 2B, Supplementary Fig. 3B and D). Glun2B protein levels in homogenates and synaptosome fractions were reduced by approximately 50% in adult heterozygous knockout (*Grin2b^+/-^*) rats compared to wild-types (*Grin2b^+/+^*) in somatosensory cortex and hippocampus (somatosensory cortex homogenate, Two-sample unpaired t-test, *DF* = 14, *T* = −5.34, *P* = 0.000104; synaptosome, Wilcoxon rank sum test, *W* = 64, *P* = 0.00094; hippocampal homogenate, Two-sample unpaired t-test, *DF* = 14, *T* = −897, *P* < 0.00001; synaptosome, Two-sample unpaired t-test, *DF* = 14, *T* = −3.89, *P* = 0.0016; Fig. 1C and Supplementary Fig. 2C). The expression of GluN2A, GluR1 and PSD95 synaptic proteins in *Grin2b^+/-^* rats was unaffected in the somatosensory cortex (GluN2A homogenate, Two-sample unpaired t-test, *DF* = 14, *T* = −0.14, *P* = 0.89; synaptosome, Two-sample unpaired t-test, *DF* = 14, *T* = −1.52, *P* = 0.15; GluR1 homogenate, Two-sample unpaired t-test, *DF* = 14, *T* = −0.98, *P* = 0.35; synaptosome, Wilcoxon rank sum test, *W* = 28, *P* = 0.71; PSD95 homogenate, Two-sample unpaired t-test, DF = 14, *T* = −0.35, *P* = 0.73; synaptosome, Two-sample unpaired t-test, *DF* = 14, *T* = 0.59, *P* = 0.56; Fig. 1D-F) and the hippocampus (GluN2A homogenate, Two-sample unpaired t-test, *DF* = 14, *T* = −0.16, *P* = 0.88; synaptosome, Two-sample unpaired t-test, *DF* = 14, *T* = 0.48, *P* = 0.64; GluR1 homogenate, Two-sample unpaired t-test, *DF* = 14, *T* = −1.57, *P* = 0.14; synaptosome, Two-sample unpaired t-test, *DF* = 14, *T* = −0.44, *P* = 0.67; PSD95 homogenate, Two-sample unpaired t-test, *DF* = 14, *T* = 0.049, *P* = 0.96; synaptosome, Wilcoxon rank sum test, *W* = 28, *P* = 0.71; Supplementary Fig. 2D-F). Despite the reduction in GluN2B expression, *Grin2b^+/-^* animals appeared, fertile and physically indistinguishable from wild-type littermates.

**Figure 1.**
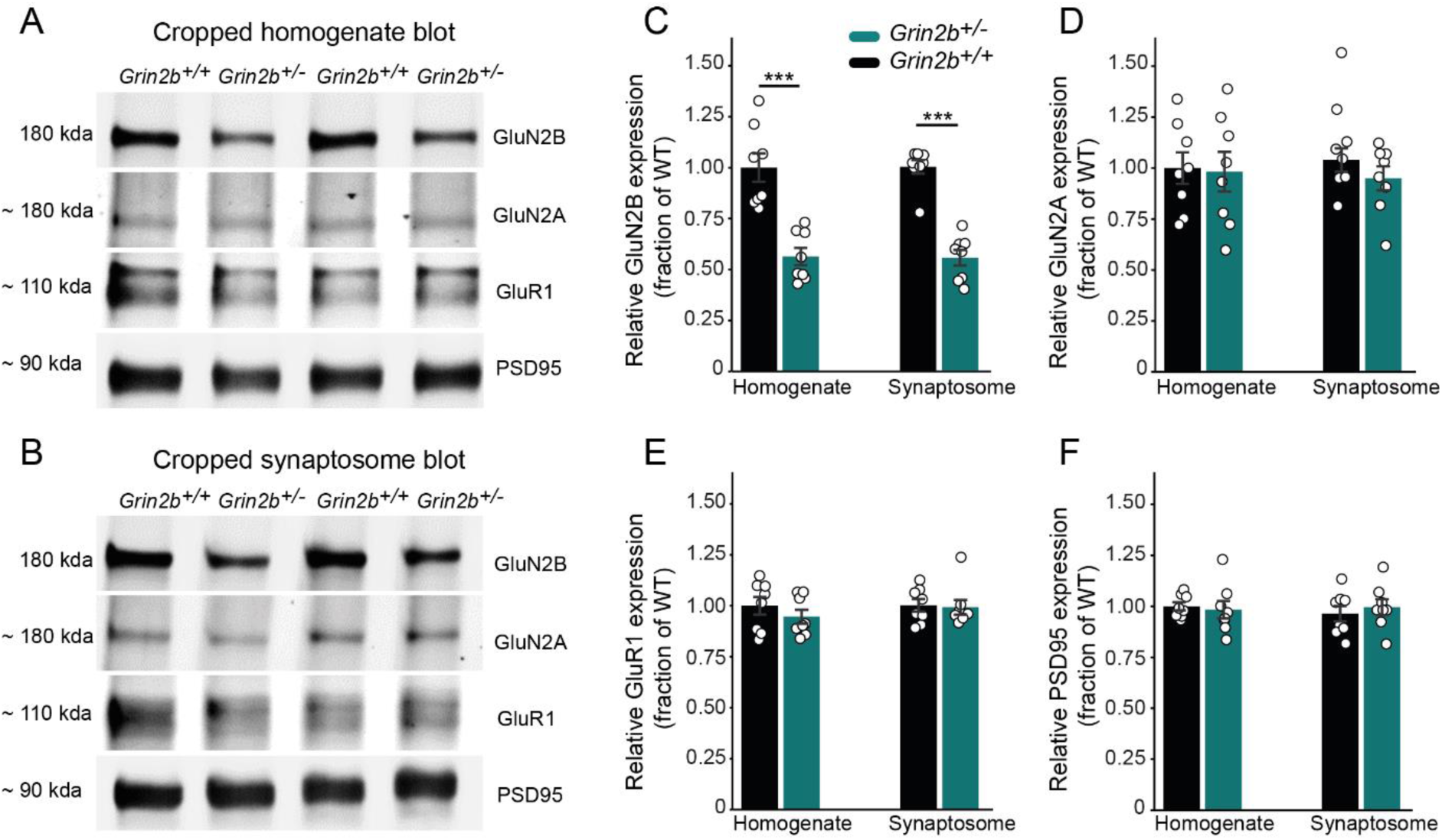
Grin2b deletion in rats results in reduction of endogenous GluN2B expression in somatosensory cortex: Representative cropped Western blots of extracts from rat somatosensory (**A**) brain homogenates and (**B**) synaptosomes. Full length blots located in Supplementary Fig 3. Bands in the molecular weight range expected for full length GluN2B, GluN2A, GluR1 and PSD95 were detected in homogenates and synaptosomes from wild-type and Grin2b^+/-^ animals. (**C**) Quantification of GluN2B protein from homogenates and synaptosomes reveals a significant decrease in Grin2b^+/-^ rats (homogenate P = 0.000104, synaptosome P = 0.00094, Two-sample unpaired t-test and Wilcoxon rank sum tests). There was no change in Grin2b^+/-^ rats in expression levels of (**D**) GluN2A (homogenate P = 0.89, synaptosome P = 0.15, Two-sample unpaired t-test), (**E**) GluR1 (homogenate P = 0.35, synaptosome P = 0.71, Two-sample unpaired t-test and Wilcoxon rank sum tests), or (**F**) PSD95 (homogenate P = 0.73, synaptosome P = 0.56, Two-sample unpaired t-test). Detailed statistics in text. Bars indicate mean values (mean ± SEM). Points correspond to values from individual rats.

### Absence seizures in *Grin2b^+/-^* rats

To determine whether heterozygous knockout of *Grin2b* results in epileptic seizures and if it affects sleep-wake brain states across the circadian cycle, animals were implanted with 14-channel surface electrode grids and EMG electrodes in neck muscles (Fig. 2A). Wireless freely moving 24-hour recordings were performed in the rats’ home cages.

*Grin2b^+/-^* rats displayed spontaneous behavioural arrests reminiscent of absence seizures which correlated with electrographic SWDs (Fig. 2A and see also Supplementary Video 1). Time-of-day factors including sleep, circadian rhythm, and light-dark phase dynamics influence seizure expression (Cho, 2012; Matos et al., 2010). We therefore first looked at the distribution of SWDs across each hour of the 24-hour cycle to evaluate whether the occurrence of SWDs in *Grin2b^+/-^*animals was higher than in wild-type littermates at specific hours of the day. We then tested whether there were any differences in the number of SWDs during the 12-hour light and dark phases, as collectively the above analyses could inform on relevant seizure activity patterns in *Grin2b^+/-^*animals. We found that there was a statistically higher number of SWDs in *Grin2b^+/-^*animals compared to controls across the entire day, and that hour of the day did not impact on this difference (Linear Mixed Model, effect of genotype *F* = 55.36, *DF* = 1, *P* < 0.00001; effect of hour *F* = 1.29, *DF* = 23*, P* = 0.16; hour x genotype *F* = 0.97, *DF* = 23 *P* = 0.49; Fig. 2B). The number of SWD events in *Grin2b^+/-^*animals did not differ between light and dark phases, and was significantly higher than that observed in wild-type rats (Linear Mixed Model, effect of genotype *F* = 55.36, *DF* = 1, *P* < 0.00001; effect of phase *F* = 1.23, *DF* = 1*, P* = 0.28; phase x genotype *F* = 0.41, *DF* = 1 *P* = 0.53; Supplementary Fig. 4A). The above reflected a statistically significant increase in the total number of SWDs over the full 24 hours in *Grin2b^+/-^*rats when compared to wild-type *Grin2b^+/+^* littermates (Wilcoxon rank sum test, *W* = 4.5, *P* = 0.0002; Fig. 2C). Furthermore, *Grin2b^+/-^* rats had a longer average SWD duration than *Grin2b^+/+^* rats (Two-sample unpaired t-test, *DF* = 22, *T* = −5.26, *P* = 0.00003; Fig. 2C) and the total duration of SWDs was also significantly longer in *Grin2b^+/-^* animals when compared to *Grin2b^+/+^* rats (Wilcoxon rank sum test, *W* = 5.5, *P* = 0.0002; Fig. 2C). These results suggest that *Grin2b* haploinsufficiency drives the network to a state of increased absence seizure activity.

**Figure 2.**
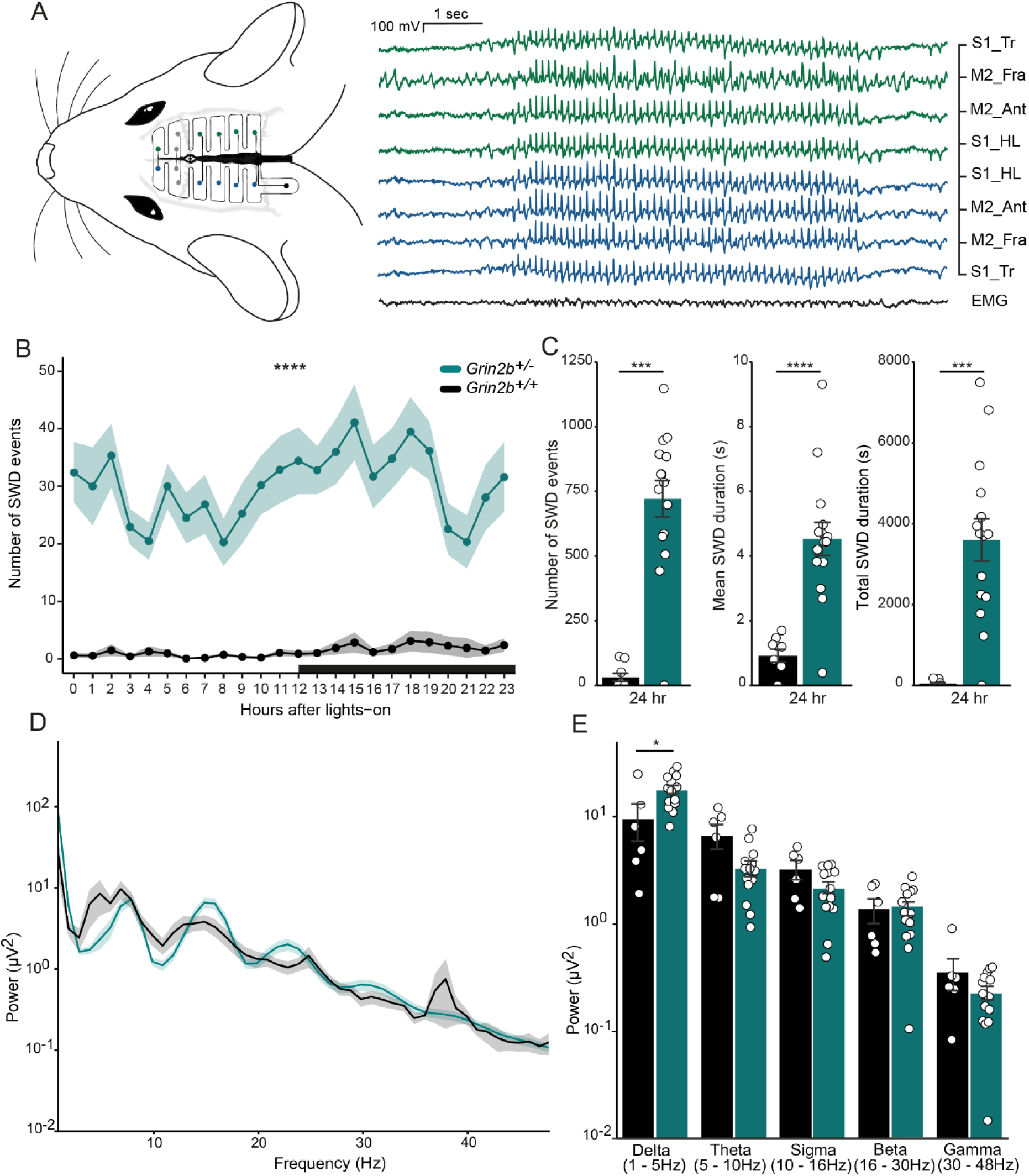
Prevalence and spectral properties of absence seizures in Grin2b^+/-^ animals: (**A**) Schematic of a 16-channel skull-surface EEG implant illustrating approximate placement of electrodes relative to the skull (left) and representative EEG and EMG voltage traces from bilateral electrode pairs in diagram showing a SWD over approximate cortical locations (S1-Tr = Primary somatosensory cortex trunk region, M2-FrA = Secondary motor cortex frontal association area, M2-Ant = Secondary motor cortex anterior, S1-HL = Primary somatosensory cortex hindlimb region) (right). (**B**) Average number of SWD events across all rats plotted by hour over the 24-hour light-dark cycle with black bar on x-axis indicating lights off in the animal facility. Hour of the day had no effect on the number of seizures between the two genotypes and Grin2b^+/-^ rats had significantly more seizures overall when compared to wild-type littermates (genotype P < 0.00001, hour P = 0.16, hour x genotype P = 0.49, Linear Mixed Model), (**** = effect from genotype). Points indicate mean values for all animals (mean ± SEM). (**C**) Bar plots of total number of SWD events (left), SWD average duration (middle) and total SWD durations (right) average across animals over the entire 24 hours with Grin2b^+/-^ rats having significantly higher values (P = 0.0002, P = 0.00003, P = 0.0002, Two-sample unpaired t-test and Wilcoxon rank sum tests). (**D**) Average EEG power spectra for Grin2b^+/-^ rats and wild-type littermates during all SWD epochs across all animals in each group. Lines represent averages for all animals (mean ± SEM). (**E**) Quantification of average power in commonly used frequency bands during SWDs, showing an increased delta power (genotype P = 0.47, frequency P < 0.00001, genotype x frequency P = 0.015, delta P = 0.039, Two-way ANOVA with Tukey post hoc tests), (* = post hoc test for interaction effect). Detailed statistics in text. Bars indicate mean values (mean ± SEM). Points correspond to values from individual rats in **C**-**E**.

We performed spectral analysis on SWDs to determine if these differed between genotypes. In SWD epoch averages we found a clear peak in the theta frequency range with a secondary harmonic in both genotypes (Fig. 2D). Spectral power during SWDs in *Grin2b^+/-^* rats was higher in the delta frequency range when compared to wild-type littermates (Two-way ANOVA, effect of genotype *F* = 0.53, *DF* = 1, *P* = 0.47; frequency *F* = 72.03, *DF* = 4, *P* < 0.00001; genotype x frequency *F* = 3.27, *DF* = 4, *P* = 0.015; Tukey *post hoc* test, delta *P* = 0.039, theta *P* = 0.34, sigma *P* = 0.704, beta *P* = 0.99, gamma *P* = 0.69; Fig. 2E). The overall increased delta power in *Grin2b^+/-^* rats may indicate that the mechanisms underlying SWD generation may favour brain states associated with heightened delta activity, such as NREM sleep.

To determine whether male and female *Grin2b^+/-^*rats have different susceptibility to the occurrence of SWDs, we compared both sexes of each genotype from our animal cohort discussed above. We found that neither sex nor hour of the day accounted for the increase in SWD events between *Grin2b^+/-^* and *Grin2b^+/+^* rats, and that differences were specific to genotype (Linear Mixed Model, effect of hour *F* = 1.24, *DF* = 23, *P* = 0.21; genotype *F* = 47.83, *DF* = 1, *P* < 0.00001; sex *F* = 0.12, *DF* = 1, *P* = 0.73, hour x genotype x sex *F* = 0.85, *DF* = 23, *P* = 0.66; Supplementary Fig. 5A).

Similarly, while there were still statistically significant differences between genotypes, there were no differences between male and female *Grin2b^+/-^* rats in the total number of SWD events (Two-way ANOVA, effect of sex *F* = 0.12, *DF* = 1, *P* = 0.74; genotype *F* = 47.663 *DF* = 1, *P* < 0.0001; sex x genotype *F* = 0.2, *DF* = 1, *P* = 0.66; Supplementary Fig. 5B), the mean duration of SWDs (Two-way ANOVA, effect of sex *F* = 0.0002, *DF* = 1, *P* = 0.99; genotype *F* = 23.17, *DF* = 1, *P* = 0.0001; sex x genotype *F* = 0.017, *DF* = 1, *P* = 0.89; Supplementary Fig. 5B) or the total time spent in a state of SWDs (Two-way ANOVA, effect of sex *F* = 0.0001, *DF* = 1, *P* = 0.99; genotype *F* = 22.9, *DF* = 1, *P* = 0.0001; sex x genotype *F* = 0.002, *DF* = 1, *P* = 0.96; Supplementary Fig. 5B).

### Sleep-wake abnormalities in *Grin2b^+/-^* rats

We quantified minutes spent in REM, NREM and wake (Fig. 3A-C) to determine if *Grin2b^+/-^* animals exhibited abnormal sleep-wake patterns as reported in patients (Freunscht et al., 2013; Simons Searchlight, 2021). Similarly to the above we tested whether *Grin2b* heterozygous knockout affects the circadian dynamics of brain states. We compared hourly and light-dark phase distributions of sleep-wake states between *Grin2b^+/-^*and *Grin2b^+/+^* rats to identify potentially dysregulated sleep-wake rhythms during specific times of the day.

**Figure 3.**
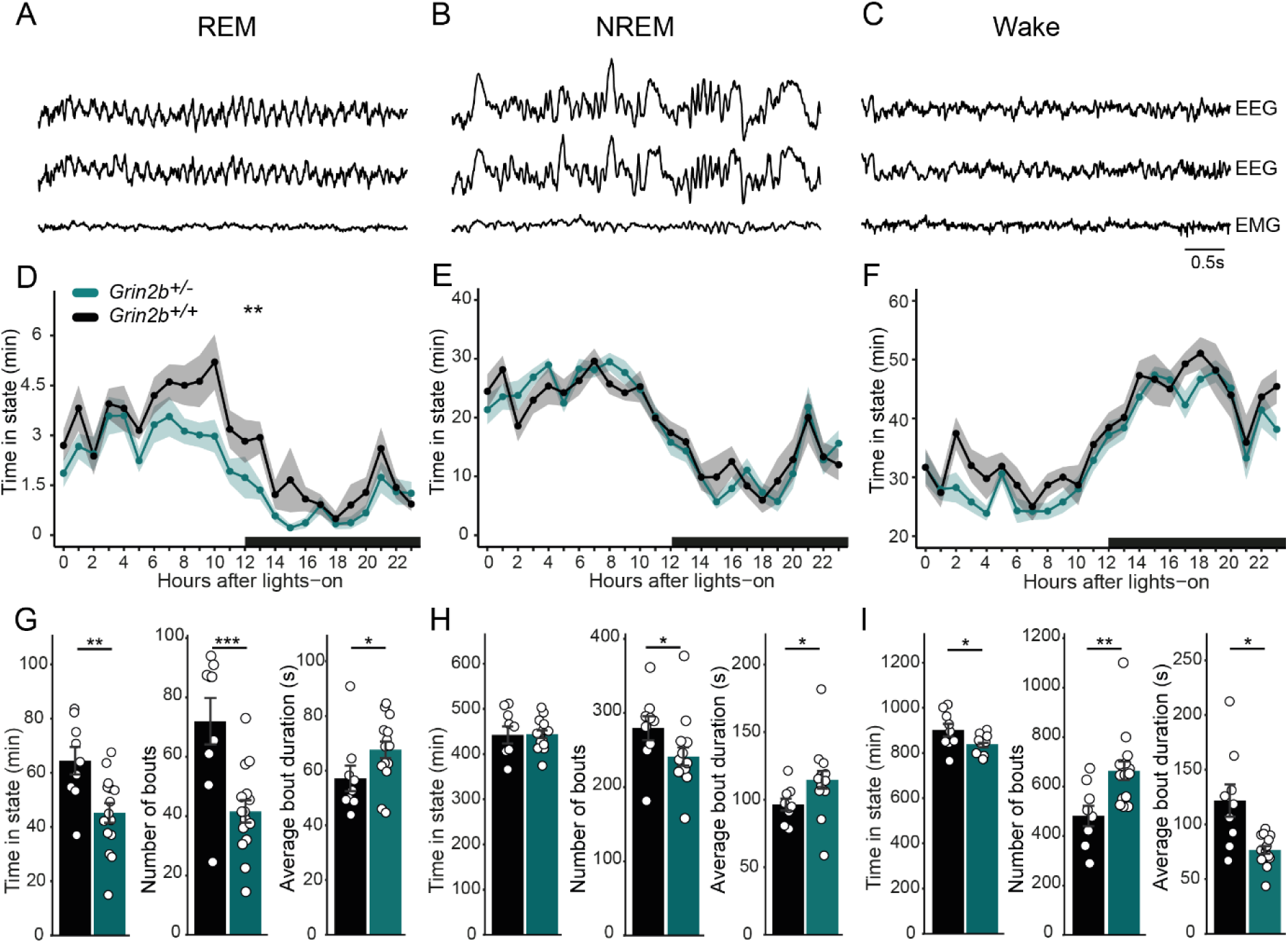
Grin2b^+/-^ animals display sleep-wake abnormalities: Representative EEG and EMG voltage traces for (**A**) REM, (**B**) NREM and (**C**) wake. Quantification showing the average time spent by hour across all rats in (**D**) REM, (**E**) NREM and (**F**) wake by hour, across the 24-hour light-dark cycle with black bar on x-axis indicating lights off in the animal facility. Grin2b^+/-^ rats had overall significantly less REM sleep compared to wild-type littermates, and this reduction was not specific to certain hours of the day (REM hour P < 0.00001, genotype P = 0.0045, hour x genotype P = 0.53; Linear Mixed Model). No net or individual hour differences were observed between genotypes for NREM and wake states (NREM hour P < 0.00001, genotype P = 0.93, hour x genotype P = 0.66; wake hour P < 0.00001, genotype P = 0.014, hour x genotype P = 0.84; Linear Mixed Model). ** = effect from genotype. Points indicate mean values for all animals (mean ± SEM). Bar plots of averages across animals showing total time (left), number of bouts (middle) and average bout duration (right) for (**G**) REM, (**H**) NREM and (**I**) wake during the full 24 hours (* = P < 0.05, ** = P < 0.01, *** = P < 0.001, unpaired Two-sample t-tests and Wilcoxon rank sum tests specified in results text). Bars indicate mean values (mean ± SEM). Points correspond to values from individual rats.

*Grin2b^+/-^* rats displayed a statistically significant net decrease in REM sleep minutes over the entire 24 hours when analysed hour-by-hour relative to wild-type animals, however, this reduction was not specific to particular hours of the day (Linear Mixed Model, effect of hour *F* = 16.13, *DF* = 23, *P* < 0.00001; genotype *F* = 9.98, *DF* = 1, *P* = 0.0045; hour x genotype *F* = 0.95, *DF* = 23, *P* = 0.53; Fig. 3D). Similarly, *Grin2b^+/-^*rats spent less total time in REM sleep than wild-type littermate controls when total minutes per animal were analysed (Two-sample unpaired t-test, *DF* = 22, *T* = 3.16, *P* = 0.0045; Fig. 3G). It is likely this reduction was due to fewer REM sleep bouts in *Grin2b^+/-^* animals (Two-sample unpaired t-test, *DF* = 22, *T* = 3.94, *P* = 0.00069; Fig. 3G). Interestingly, the average duration of REM sleep bouts was longer for *Grin2b^+/-^* animals (Wilcoxon rank sum test, *W* = 33, *P* = 0.043) (Fig. 3G), however, this was insufficient to compensate for the net REM deficit observed. The significantly reduced REM sleep in *Grin2b^+/-^* mutants was not specific to light or dark periods (Linear Mixed Model, effect of phase *F* = 119.42, *DF* = 1, *P* < 0.00001; genotype *F* = 9.99, *DF* = 1, *P* = 0.0046; phase x genotype *F* = 0.85, *DF* = 1, *P* = 0.37; Supplementary Fig. 6A). We did however, observe significantly fewer REM sleep bouts during both the light and dark periods in *Grin2b^+/-^* rats (Linear Mixed Model, effect of phase *F* = 105.93, *DF* = 1, *P* < 0.00001; genotype *F* = 15.52, *DF* = 1, *P* = 0.00067; phase x genotype *F* = 4.53, *DF* = 1, *P* = 0.045; Tukey *post hoc* test, light *P* = 0.0001, dark *P* = 0.032; Supplementary Fig. 6A). The average duration of REM sleep bouts did not significantly differ between the light and dark phase between genotypes (Linear Mixed Model, effect of phase *F* = 2.26, *DF* = 1, *P* = 0.13; genotype *F* = 1.88, *DF* = 1, *P* = 0.18; phase x genotype *F* = 2.15, *DF* = 1, *P* = 0.16; Supplementary Fig. 6A).

On an hour-by-hour basis the distribution of NREM sleep was not different between *Grin2b^+/-^* and *Grin2b^+/+^* rats (Linear Mixed Model, effect of hour *F* = 21.52, *DF* = 23, *P* < 0.00001; genotype *F* = 0.0081, *DF* = 1, *P* = 0.93; hour x genotype *F* = 0.86, *DF* = 23, *P* = 0.66; Fig. 3E). Similarly, there was no difference between *Grin2b^+/-^* and *Grin2b^+/+^* rats in the total amount of time spent in NREM over the full 24 hours (Two-sample unpaired t-test, *DF* = 22, *T* = −0.09, *P* = 0.93; Fig. 3H). Nonetheless, *Grin2b^+/-^* rats demonstrated abnormal NREM distributions when compared to *Grin2b^+/+^* littermates, as they had statistically fewer NREM bouts (Wilcoxon rank sum test, *W* = 29.5, *P* = 0.025; Fig. 3H), which where longer in average duration (Wilcoxon rank sum test, *W* = 26, *P* = 0.015; Fig. 3H). During the light and dark phases, however, *Grin2b^+/-^* did not differ from their wild-type littermates in the total amount of NREM sleep (Linear Mixed Model, effect of phase *F* = 237.34, *DF* = 1, *P* < 0.00001; genotype *F* = 0.0065, *DF* = 1, *P* = 0.94; phase x genotype *F* = 0.94, *DF* = 1, *P* = 0.34; Supplementary Fig. 6B), the amount of NREM bouts (Linear Mixed Model, effect of phase *F* = 198.28, *DF* = 1, *P* < 0.00001; genotype *F* = 3.62, *DF* = 1, *P* = 0.07; phase x genotype *F* = 0.041, *DF* = 1, *P* = 0.84; Supplementary Fig. 6B) and the average NREM bout duration (Linear Mixed Model, effect of phase *F* = 16.34, *DF* = 1, *P* = 0.00055; genotype *F* = 4.091, *DF* = 1, *P* = 0.055; phase x genotype *F* = 0.0023, *DF* = 1, *P* = 0.99; Supplementary Fig. 6B).

Finally, we evaluated the distribution of wake across each hour of the light-dark cycle and found that while overall *Grin2b^+/-^*animals spent significantly reduced time awake, this was not specific to any given hour (Linear Mixed Model, effect of hour *F* = 22.55, *DF* = 23, *P* < 0.00001; genotype *F* = 7.14, *DF* = 1, *P* = 0.014; hour x genotype *F* = 0.71, *DF* = 23, *P* = 0.84; Fig. 3F). When compared to wild-type littermates, the total minutes awake were significantly reduced in *Grin2b^+/-^* rats (Welch’s two-sample unpaired t-test, *DF* = 10.25, *T* = 2.26, *P* = 0.047; Fig. 3I) with mutants having more wake bouts (Wilcoxon rank sum test, *W* = 18, *P* = 0.0035; Fig. 3I) of shorter duration (Welch’s two-sample unpaired t-test, *DF* = 8.91, *T* = 2.97, *P* = 0.016; Fig. 3I). Wake state differences were due to the overall changes across the full 24 hours, as the differences between *Grin2b^+/-^* and *Grin2b^+/+^*animals were irrespective of the light or dark period for the time spent awake (Linear Mixed Model, effect of phase *F* = 185.12, *DF* = 1, *P* < 0.00001; genotype *F* = 6.64, *DF* = 1, *P* = 0.013; phase x genotype *F* = 0.23, *DF* = 1, *P* = 0.64; Supplementary Fig. 6C), number of wake bouts (Linear Mixed Model, effect of phase *F* = 1.39, *DF* = 1, *P* = 0.25; genotype *F* = 9.76, *DF* = 1, *P* = 0.0049; phase x genotype *F* = 1.27, *DF* = 1, *P* = 0.27; Supplementary Fig. 6C) and average wake bout duration (Linear Mixed Model, effect of phase *F* = 26.25, *DF* = 1, *P* < 0.00001; genotype *F* = 13.14, *DF* = 1, *P* = 0.0015; phase x genotype *F* = 2.34, *DF* = 1, *P* = 0.14; Supplementary Fig. 6C).

Sleep states are altered in *Grin2b^+/-^*rats. However, there were no statistically significant differences between male and female mutant and wild-type animals (for statistical results see Supplementary Table 2) (Supplementary Fig. 7A-F).

We hypothesized that heterozygous *Grin2b* knockout may affect the physiological dynamics of individual brain states. We therefore tested whether spectral properties across REM, NREM and wake in *Grin2b^+/-^* rats differed from those of *Grin2b^+/+^* animals. We calculated the average EEG spectral power from one representative channel across all animals and epochs of each brain state. No statistical differences were found between *Grin2b^+/-^* and *Grin2b^+/+^* rats during REM sleep (Two-way ANOVA, effect of genotype *F* = 2.25, *DF* = 1, *P* = 0.14; frequency *F* = 281.56, *DF* = 4, *P* < 0.00001; genotype x frequency *F* = 0.57, *DF* = 4, *P* = 0.69; Fig. 4A). However, significant differences in spectral power were found for wake and NREM sleep. In NREM sleep, *Grin2b^+/-^*animals have statistically higher beta power in comparison to wild type rats (Two-way ANOVA, effect of genotype *F* = 5.89, *DF* = 1, *P* = 0.017; frequency *F* = 4037.23, *DF* = 4, *P* < 0.00001; genotype x frequency *F* = 2.64, *DF* = 4, *P* = 0.038; Tukey *post hoc* test, delta *P* = 0.98, theta *P* = 0.99, sigma *P* = 0.82, beta *P* = 0.0008, gamma *P* = 0.99; Fig. 4B). While in the wake state spectral power was significantly reduced in *Grin2b^+/-^* rats relative to *Grin2b^+/+^* littermates, although this reduction was not specific to individual frequency bands (Two-way ANOVA, effect of genotype *F* = 6.84, *DF* = 1, *P* = 0.01; frequency *F* = 657.61, *DF* = 4, *P* < 0.00001; genotype x frequency *F* = 0.33, *DF* = 4, *P* = 0.86, Fig. 4C). Grouped together, the above results show that *Grin2b^+/-^* rats have altered sleep-wake brain state distributions and spectral properties.

**Figure 4.**
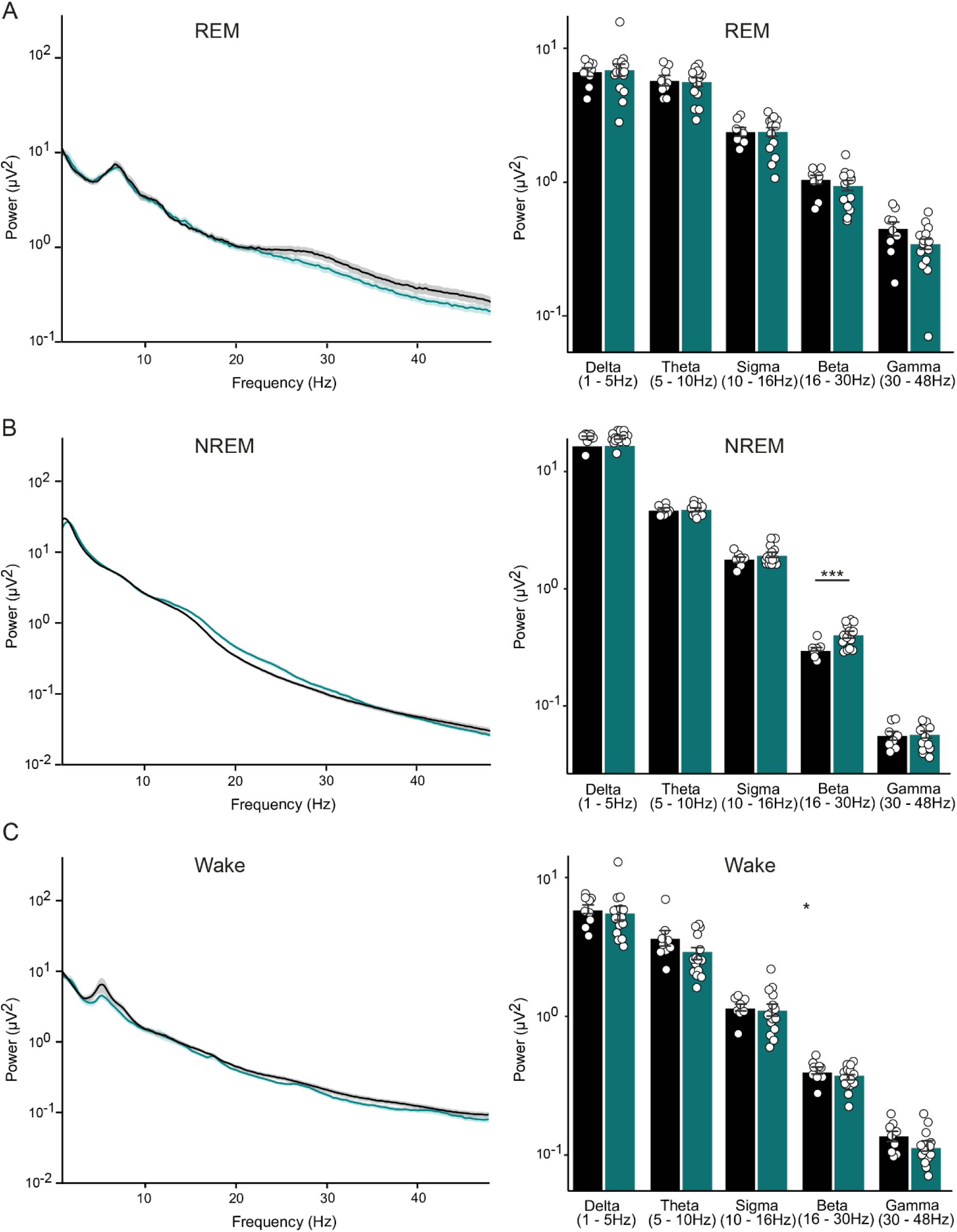
Spectral power is altered in wake and NREM sleep in Grin2b^+/-^ rats: Average EEG power spectra across wild-type and Grin2b^+/-^ rats (left; traces represent averages for all animals (mean ± SEM)) and quantification by individual frequency bands (right) during (**A**) REM, (**B**) NREM and (**C**) wake epochs. Quantification of average power in specific frequency bands during REM epochs showed no spectral differences between Grin2b^+/-^ and wild-type animals (genotype P = 0.14, frequency P < 0.00001, genotype x frequency P = 0.69; Two-way ANOVA). Spectral power in the beta frequency range was significantly higher during NREM epochs in Grin2b^+/-^ rats (genotype P = 0.017, frequency P < 0.00001, genotype x frequency P = 0.038 (beta P = 0.0008), Two-way ANOVA with Tukey post hoc tests), (*** = post hoc test for interaction). During wake, in Grin2b^+/-^ animals overall power was reduced relative to Grin2b^+/+^ littermates, although no significant differences were found in individual frequency bands (genotype P = 0.01, frequency P < 0.00001, genotype x frequency P = 0.86; Two-way ANOVA), (* = effect from genotype). Detailed statistics in main text. Bars indicate mean values (mean ± SEM). Points correspond to values from individual rats.

### Brain state abnormalities in *Grin2b^+/-^* rats are not correlated with absence seizure activity

We hypothesized that the increased SWD activity in *Grin2b^+/-^* rats may contribute to the altered sleep-wake brain states. We therefore first tested whether a higher percentage of SWDs preferentially occurred during wake, NREM or REM sleep and whether there were any differences in this observation between *Grin2b^+/-^* and *Grin2b^+/+^* rats. Collectively for both genotypes, we observed that SWDs predominantly occurred during wake (94.8%), followed by NREM (4.4%) and scarcely during REM sleep (0.5%) (Fig. 5A). Rats have polyphasic sleep patterns, with wake bouts occurring throughout the light and dark phase periods of the 24-hour cycle (Yasenkov & Deboer, 2012), and these data further highlight that SWD occurrence is more likely state than light or dark phase dependent. Interestingly, a higher percentage of SWDs initiated during NREM in *Grin2b^+/-^* rats than in wild-type littermates, while a higher percentage of seizures in wild-type animal occurred during the wake state (Two-way ANOVA, effect of genotype *F* = 1, *DF* = 1, *P* = 0.98; state *F* = 5945.056, *DF* = 2, *P* < 0.00001; genotype x state *F* = 10.94, *DF* = 2, *P* = 0.00008; Tukey *post hoc* test, wake *P* = 0.0022, NREM *P* = 0.0011; Fig. 5A). The increased frequency of *Grin2b^+/-^* SWDs in NREM sleep prompted us to examine whether the wake-NREM transitional state (Fig. 5B and C), when synchronous subcortical output is increased (Halassa et al., 2014), is linked to SWD occurrence and if this differed between groups. We tested all SWD epochs identified to originate in NREM and found that all *Grin2b^+/-^*rats had SWDs initiating at wake-NREM transitions, while this was the case for only a small fraction of *Grin2b^+/+^*littermates (33.33 %, 3/9 rats) (Fishers exact test, *P* = 0.00062; Fig. 5C). We subsequently evaluated for differences in the proportion of wake-NREM transitional SWDs between *Grin2b^+/-^* animals and wild-type rats that presented with transitional state seizures. However, we did not observe any variation in the percent of wake-NREM transitional SWDs between the two groups, 24.2% of SWDS in *Grin2b^+/-^* rats occurred during the wake-NREM transitional period, which was similar to the 22.39% observed in *Grin2b^+/-^* littermates (Two-sample unpaired t-test, *DF* = 16, *T* = −0.34, *P* = 0.74; Supplementary Fig. 4B).

**Figure 5.**
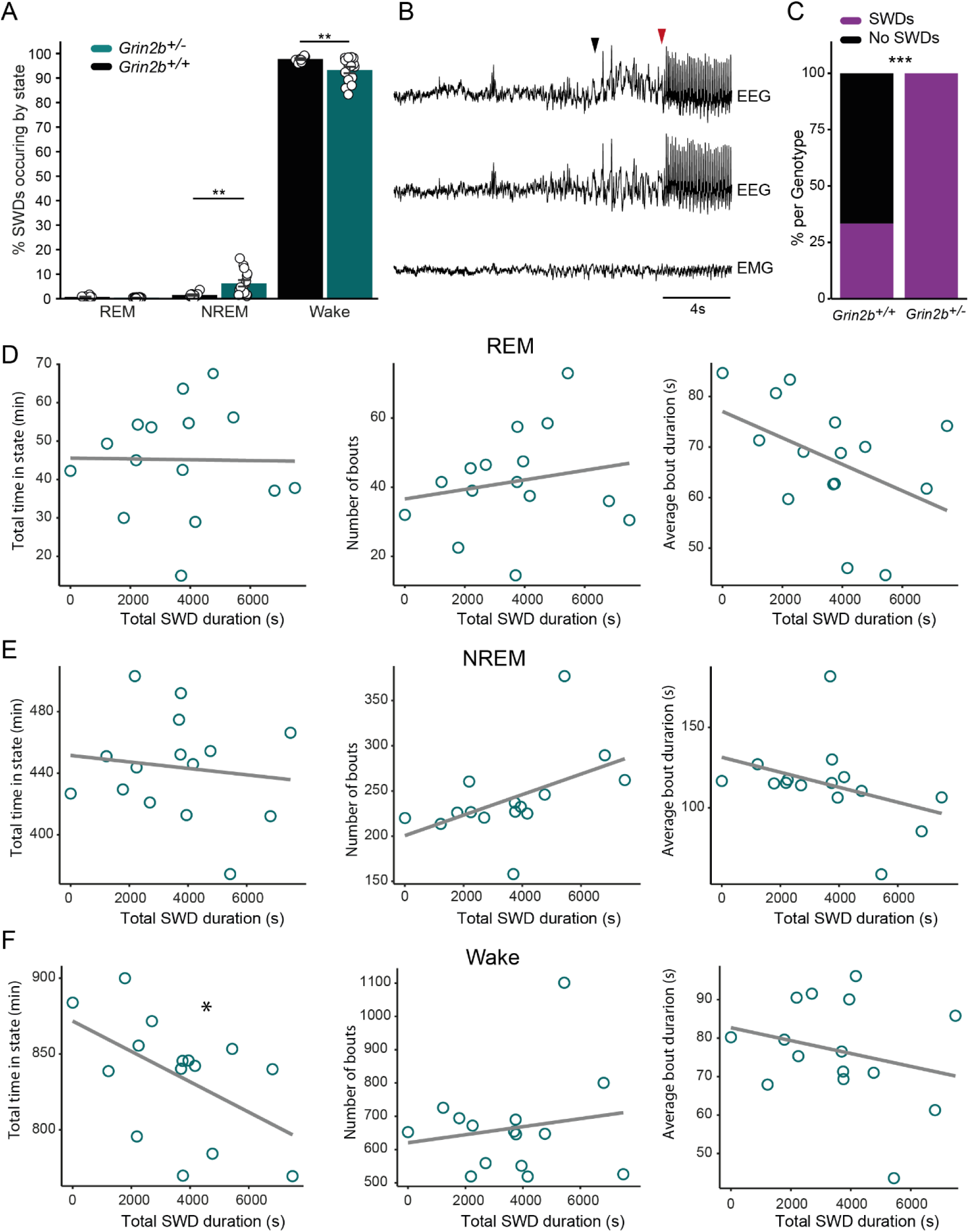
SWDs preferentially occur during wake but sleep abnormalities do not correlate with SWDs in Grin2b^+/-^ animals: (**A**) Plot of average percentage of SWDs occurring during REM, NREM and wake brain states. Compared to wild-type littermates, Grin2b^+/-^ rats have less SWDs in wake and more SWDs in NREM (genotype P = 0.98, state P < 0.00001, genotype x state P = 0.00008 (wake P = 0.0022, NREM P = 0.0011), Two-way ANOVA with Tukey post hoc tests). Bars indicate mean values (mean ± SEM). Points correspond to values from individual rats. (**B**) Representative EEG and EMG voltage traces during a wake-NREM transition with a SWD. Black arrowhead indicates start of NREM and red arrowhead indicates start of SWD. (**C**) A higher proportion of SWDs in NREM initiate during wake-NREM transitions in Grin2b^+/-^ when compared to wild-type animals (P = 0.00062, Fishers exact test). Bars indicate all animals per each genotype and fill colouring indicates the relevant proportion of animals with SWDs. Points correspond to values from individual rats. Correlation plots of total time in state (left), number of bouts (middle) and average bout duration (right) with SWD duration for (**D**) REM, (**E**) NREM and (**F**) wake, plotted against total SWD duration. Quantification revealed total SWD duration is negatively correlated to total time awake (P = 0.048, Pearson correlation) and is uncorrelated to all other sleep measurements (see results section for detailed statistics). Points correspond to values from individual Grin2b^+/-^ rats.

We hypothesized that there might be a mechanistic link between sleep-wake abnormalities and SWD occurrence. We therefore next determined whether there was a correlation between sleep-wake properties and total SWD duration in *Grin2b^+/-^*rats. Total seizure duration in *Grin2b^+/-^* animals did not correlate with total time in REM (Pearson correlation, *T* = −0.055, *DF* = 13, *R* = - 0.015, *P* = 0.96; Fig. 5D) and NREM sleep (Pearson correlation, *T* = −1.47, *DF* = 13, *R* = −0.13, *P* = 0.65; Fig. 5E). There was also no correlation between total SWD duration and bouts of REM (Pearson correlation, *T* = 0.704, *DF* = 13, *R* = −0.19, *P* = 0.49; Fig. 5D) or NREM (Pearson correlation, *T* = 2.006, *DF* = 13, *R* = 0.49, *P* = 0.066; Fig. 5E), as well as average bout duration of REM (Pearson correlation, *T* = 0.61, *DF* = 13, *R* = 0.17, *P* = 0.55; Fig. 5D), or NREM (Pearson correlation, *T* = −1.44, *DF* = 13, *R* = −0.37, *P* = 0.17; Fig. 5E). Finally, we found a correlation between *Grin2b^+/-^* total SWD durations and total time in wake (Pearson correlation, *T* = −2.16, *DF* = 13, *R* = −0.51, *P* = 0.048; Fig 5F). This is possibly linked to the statistically decreased time in wake in *Grin2b^+/-^*rats (Fig. 3I), which have much higher numbers of SWDs than controls (Fig. 2C). In other words, in *Grin2b^+/-^* animals, SWDs likely replace time in the wake state. There was no correlation, however, between total duration of seizures with wake bouts (Pearson correlation, *T* = 0.61, *DF* = 13, *R* = 0.17, *P* = 0.55; Fig. 5F) or average wake bout duration (Pearson correlation, *T* = −0.93, *DF* = 13, *R* = −0.25, *P* = 0.37; Fig. 5F) in *Grin2b^+/-^* mutants. The general lack of correlation between SWDs and sleep properties in *Grin2b^+/-^*animals suggests that the two phenotypes have independent underlying mechanisms.

### Pharmacological blockade of SWDs

We next evaluated whether SWDs in *Grin2b^+/-^*and wild-type rats could be suppressed pharmacologically. We tested the efficacy of ethosuximide, a T-type voltage-gated calcium channel blocker used in the treatment of absence seizures in humans (Zimmerman & Burgemeister, 1958) and which is known to block SWDs in other absence seizure rodent models (Terzioglu et al., 2006). We also evaluated seizure responses to memantine, a noncompetitive NMDAR antagonist currently explored as a mono or adjunctive treatment option in NMDAR related epilepsies (Bormann, 1989; H. S. Chen & Lipton, 1997; Parsons et al., 1999; Platzer et al., 2017). As *Grin2b^+/-^* rats had a more severe seizure phenotype, we hypothesized that altered NMDAR-mediated signalling may contribute to brain circuits underlying SWDs, and that blocking NMDARs would therefore reduce SWD activity.

We found that in both *Grin2b^+/-^* and *Grin2b^+/+^* rats there was a statistically significant reduction in SWDs with pharmacological treatment, as systemic ethosuximide resulted in significantly fewer SWDs than memantine and saline (Linear Mixed Model, effect of treatment *F* = 6.48, *DF* = 2, *P* = 0.0049; genotype *F* = 0.42, *DF* = 1, *P* = 0.53; treatment x genotype *F* = 0.34, *DF* = 2, *P* = 0.71; Tukey *post hoc* test for effect of treatment, ethosuximide-saline *P* = 0.0081, ethosuximide-memantine *P* = 0.015,, memantine-saline *P* = 0.96; Fig. 6B). Both ethosuximide and memantine, however, reduced the average duration of SWDs in *Grin2b^+/-^* but not *Grin2b^+/+^* animals (Linear Mixed Model, effect of treatment *F* = 9.17, *DF* = 2, *P* = 0.00048; genotype *F* = 0.71, *DF* = 1, *P* = 0.41; treatment x genotype *F* = 3.34, *DF* = 2, *P* = 0.044; Tukey *post hoc* test for interaction effect by genotype, *Grin2b^+/-^* ethosuximide-saline *P* < 0.0001, *Grin2b^+/-^* memantine-saline *P* = 0.0009, *Grin2b^+/-^* memantine-ethosuximide *P* = 0.76, *Grin2b^+/+^* ethosuximide-saline *P* = 0.88, *Grin2b^+/+^* memantine-saline *P* = 0.99, *Grin2b^+/+^*memantine-ethosuximide *P* = 0.99; Fig. 6C). The specific effect on *Grin2b^+/-^* rats, but not wild-types, may stem from the significantly fewer and shorter seizures observed in *Grin2b^+/+^* animals. In contrast, the effect on *Grin2b^+/-^* rats, but not wild-types, could be dependent on distinct mechanisms underlying seizures between genotypes. Furthermore, memantine’s effect on reducing SWD duration, but not affecting SWD occurrence, may be due to its use-dependent mechanism of action, requiring opening of the channel to get access to the channel pore, which only happens after seizures begin (Parsons et al., 1993). The combined effect of less SWDs with lower duration likely resulted in ethosuximide, but not memantine, reducing total seizure durations in both rat groups (Linear Mixed Model, effect of treatment *F* = 9.44, *DF* = 2, *P* = 0.0011; genotype *F* = 0.67, *DF* = 1, *P* = 0.43; treatment x genotype *F* = 1.15, *DF* = 2, *P* = 0.34; Tukey *post hoc* test for effect of treatment, ethosuximide-saline *P* = 0.0005, ethosuximide-memantine *P* = 0.083, memantine-saline *P* = 0.098; Fig. 6D). Both ethosuximide and memantine could therefore potentially be tested as treatments for SWDs in *GRIN2B* patients.

**Figure 6.**
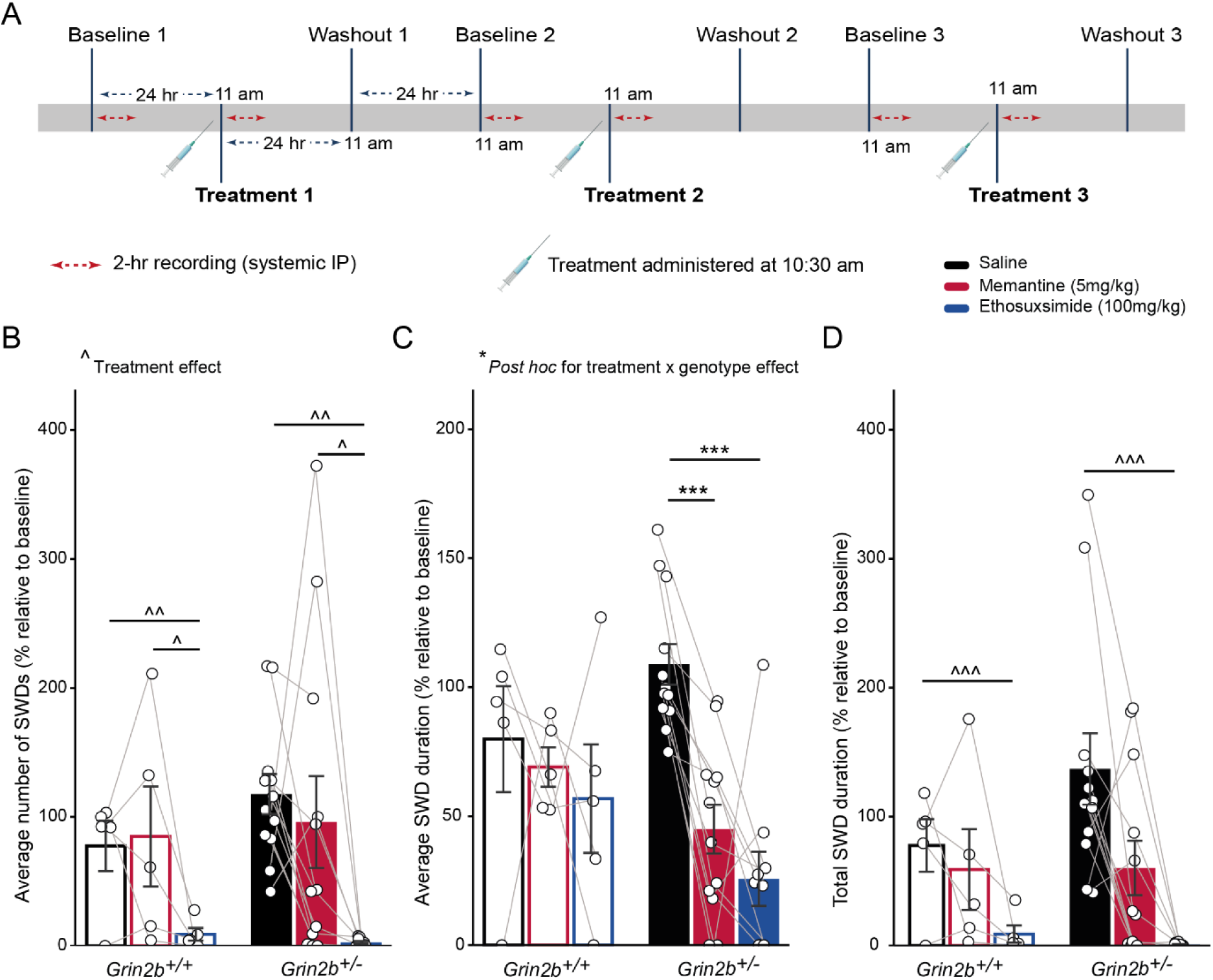
Acute administration of anti-seizure drugs attenuates SWDs: (**A**) Diagram of pharmacology experiment timeline. Note: Treatment order is randomized and weighted across animals to ensure order permutations occur equally. Plots showing effects of acute drug treatment on SWD percent relative to baseline (**B**) average number, (**C**) average duration and (**D**) total duration in Grin2b^+/-^ and wild-type animals. Treatment with ethosuximide successfully blocked SWD events in both groups (treatment P = 0.0049 (ethosuximide-saline P = 0.0081, ethosuximide-memantine P = 0.015), genotype P = 0.53, treatment x genotype P = 0.71, Linear Mixed Model with Tukey post hoc tests, ^ and ^^ = post hoc tests for treatment main effect). Memantine and ethosuximide reduced average SWD duration in Grin2b^+/-^ but not Grin2b^+/+^ rats (treatment P = 0.00048, genotype P = 0.41, treatment x genotype P = 0.044 (Grin2b^+/-^ ethosuximide-saline P < 0.0001, Grin2b^+/-^ memantine-saline P = 0.0009), Linear Mixed Model with Tukey post hoc tests, *** = post hoc test for treatment x genotype interaction effect). Total seizure duration was also significantly reduced following ethosuximide treatment in both Grin2b^+/-^ and wild-type rats (treatment P = 0.0011 (ethosuximide-saline P = 0.0005), genotype P = 0.43, treatment x genotype P = 0.34, Linear Mixed Model with Tukey post hoc tests, ^^^ = post hoc test for treatment main effect). See results for detailed statistics. Bars indicate mean values (mean ± SEM). Points correspond to values from individual rats and grey lines follow treatment response of each individual rat.

Previous research demonstrates that inhibiting T-type calcium channels in nRT effectively reduces SWDs in Genetic Absence Epilepsy Rats from Strasbourg (GAERS) (McCafferty et al., 2018; Richards et al., 2003). Additionally, in GAERs, thalamic NMDAR signalling plays a key role in regulating absence seizures of genetic origin, as both NMDAR agonists and antagonists suppress SWD activity (Koerner et al., 1996). Therefore, we next investigated the effect of ethosuximide and memantine applied directly to the nRT of Grin2b^+/-^ animals. The experimental timeline is illustrated in Fig. 7A. Infusion of ethosuximide and memantine into the nRT of Grin2b^+/-^ animals lead to a significant decrease in the amount of SWDs compared to saline (Linear Mixed Model, effect of treatment F = 31.35, DF = 2, P = 0.00067; Tukey post hoc test, ethosuximide-saline P = 0.0016, ethosuximide-memantine P = 0.75, memantine-saline P = 0.0009; Fig 7B and C). Relative to treatment with saline however, ethosuximide and memantine had no effect on reducing average SWD durations of seizures that did occur (Linear Mixed Model, effect of treatment F = 1.41, DF = 2, P = 0.32; Fig. 7B and C). Ultimately, the strong reduction in SWD events following ethosuximide and memantine administration to the nRT, resulted in decreased total seizure durations in Grin2b^+/-^ rats, when treated with either drug compared to saline control (Linear Mixed Model, effect of treatment F = 31.87, DF = 2, P = 0.00064; Tukey post hoc test, ethosuximide-saline P = 0.0016, ethosuximide-memantine P = 0.68, memantine-saline P = 0.0008; Fig. 7B and C). Overall, these data suggest that although both ethosuximide and memantine were able to modulate nRT activity to reduce SWD, likely by inhibiting nRT burst firing (McCafferty et al., 2018), once seizures initiated within the thalamocortical circuit, the network dynamics sustaining SWD activity remain largely unaffected.

**Figure 7.**
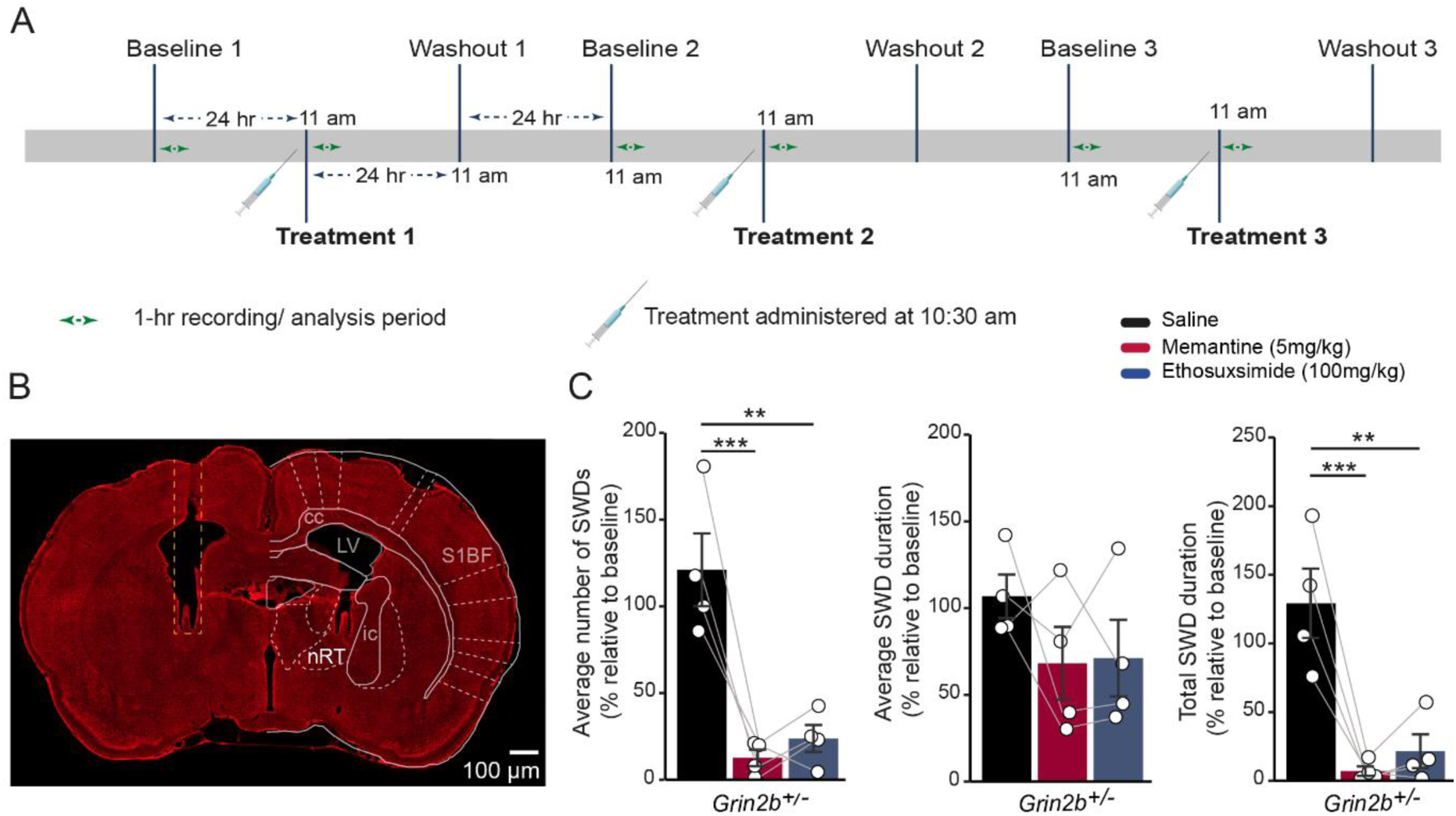
Site-specific infusion of anti-seizure drugs in nRT attenuates SWDs in Grin2b^+/-^ rats: (**A**) Diagram of pharmacology experiment timeline. Note: Treatments order is randomized and weighted across animals to ensure order permutations occur equally. (**B**) Representative histology image showing bilaterally implanted cannulas in the rostral nRT (yellow dashed lines), overlaid with approximate coordinates from Paxinos and Watson’s Rat Brain Atlas (Paxinos & Watson, 2018). S1BF = Primary somatosensory cortex barrel field, nRT = Reticular thalamic nucleus, LV = Lateral ventricle, cc = Corpus callosum, ic = Internal capsule. For detailed statistics see results section. (**C**) Panels showing effects of acute nRT drug infusion on SWD percent relative to baseline average number (left), average duration (middle), and total duration (right) of SWDs in Grin2b^+/-^ rats. Treatment with ethosuximide and memantine successfully blocked SWD events in Grin2b^+/-^ animals (treatment P = 0.00067 (ethosuximide-saline P = 0.0016, memantine-saline P = 0.0009), Linear Mixed Model with Tukey post hoc tests), (** and *** = post hoc tests for treatment effect). Neither drug reduced average SWD durations in Grin2b^+/-^ rats (treatment P = 0.32, Linear Mixed Model). Total seizure duration was also significantly reduced following ethosuximide and memantine infusion to the nRT (treatment P = 0.00064 (ethosuximide-saline P = 0.0016, memantine-saline P = 0.0008), Linear Mixed Model with Tukey post hoc tests, ** and *** = post hoc tests for treatment effect). Bars indicate mean values (mean ± SEM). Points correspond to values from individual Grin2b^+/-^ rats and grey lines follow treatment response of each individual Grin2b^+/-^ rat.

## Discussion

We identify clinically relevant epileptic and sleep-wake phenotypic abnormalities in a novel model of *GRIN2B* haploinsufficiency. *Grin2b^+/-^* animals had a 50 % reduction in *Grin2b* expression compared to wild-type levels both in the somatosensory cortex and the hippocampus in adult rats. We show that *Grin2b^+/-^* rats displayed a higher prevalence of absence seizure SWDs, which had increased delta spectral power, throughout both light and dark phases of the 24-hour cycle. Sleep-wake distributions are disrupted in mutant animals, although the occurrence of SWDs is largely uncorrelated to the severity of sleep-wake abnormalities. Finally, we found that pharmacological treatment with ethosuximide was able to both reduce the number and average duration of absence seizures while memantine only lowers average SWD duration. However, both drugs reduce absence seizure durations only in *Grin2b^+/-^* animals and not wild-types. The difference in seizure susceptibility and drug effects between heterozygous and wild-type animals could be due to direct effects of the mutation on seizure generating areas or on parallel brain mechanisms. Finally, we demonstrate that the thalamus is critically engaged in the generation of SWDs in *Grin2b^+/-^* animals by infusing the drugs directly into the nRT. Both memantine and ethosuximide are approved for clinical use and could prove to be translationally relevant options.

We show that *Grin2b^+/-^* rats had a higher number of absence seizures, which were significantly longer in duration and had higher spectral power when compared to wild-type littermates. Over 24 hours, *Grin2b^+/-^* mutants had on average 3598 seconds of SWDs, almost an entire hour of the day in an absence seizure. In contrast, wild-type littermates had average totals of less than a minute. Absence seizures have been reported in some individuals with *GRIN2B* missense mutations, although the functional impact of these variants is unknown (Epi4K Consortium., 2013).

A common finding associated with NDD and epilepsy phenotypes in various rodent models is NMDAR hypofunction. Diminished NMDAR activity has been shown in transfected rat cortical neurons null for *Shank-3* (Duffney et al., 2013), in *Shank-2* knockout mice that exhibit ASD-like impairments in social interactions, memory, and repetitive behaviours (Lim et al., 2017; Won et al., 2012), in *Fmr1* knockout mice with deficits in contextual learning (Bostrom et al., 2015; Eadie et al., 2012; Yau et al., 2016), and in rats with the *Grin2b-*Trp373 loss of function mutation (X. Wang et al., 2022). Notably, *GRIN2A* variants that decrease NMDAR activation are reported in individuals with developmental delay and severe epilepsy (Gao et al., 2017). This implies that similarly to the above, *Grin2b* haploinsufficiency could contribute to NMDAR hypofunction, seizure pathogenesis and sleep impairment in *Grin2b^+/-^* animals.

Overall, epilepsy in *GRIN2B* related NDDs has been challenging to model, especially as complete loss of GluN2B-NMDARs is perinatally lethal. Mice with heterozygous *Grin2b* knockout or a single copy of the *Grin2b*-C456Y amino binding domain variant exhibit behaviours reminiscent of ASD, but notably lack epileptic activity (O’Roak et al., 2011; Shin et al., 2020). Adult rats carrying the *Grin2b-*Trp373 loss of function variant have a reduced threshold to pentylenetetrazol induced seizures, however, they do not present with spontaneous epileptic phenotypes (X. Wang et al., 2022). Lethal status epilepticus was reported in juvenile mice lacking GluN2B specifically in inhibitory neurons (Kelsch et al., 2014), and while significant, this observation does not effectively translate to the pan neuronal impact of *GRIN2B* mutations in patients with epilepsy. Hence, the SWDs observed in *Grin2b^+/-^* mutants here represents the first report of spontaneously occurring seizures in a model with construct validity, highlighting its potential utility for future research into the contribution of heterozygous GluN2B loss to the development and pathophysiology of seizures.

Increased cortical excitation in layer V/VI is thought to trigger SWDs (Meeren et al., 2002; Polack et al., 2007; Pumain et al., 1992; Studer et al., 2019). Activity then spreads to thalamic structures and generalises bilaterally through thalamocortical circuits, which compromises reciprocally connected cortex and thalamic nuclei. Within the thalamus, excitatory thalamocortical relay neurons in the ventrobasal region and inhibitory cells in the nRT play crucial roles in modulating circuit oscillations, including SWDs (Crunelli et al., 2020; Lindquist et al., 2023). GluN2B containing NMDARs are expressed throughout the thalamocortical circuit and the two-fold decrease in GluN2B expression observed in *Grin2b^+/-^* rats likely alters thalamocortical synaptic dynamics and connectivity, similarly to other genetic animal models, in which synaptic transmission within the thalamocortical network is altered and results in SWDs (Barad et al., 2012; Cao et al., 2020; Makinson et al., 2017; Menuz & Nicoll, 2008; Paz et al., 2011).

GluN2B-NMDARs are the dominant postsynaptic receptor subtype facilitating intra-cortical signalling between layer V excitatory cells (Kumar & Huguenard, 2003), which also project to the relay thalamus and nRT (Deschênes et al., 1994; Hádinger et al., 2023; Paz et al., 2011; Pinault et al., 1995). Cortical SWD initiation could therefore be a result of insufficient GluN2B mediated NMDAR signalling at short range cortico-cortical synapses in *Grin2b^+/-^* animals. Furthermore, postsynaptic GluN2B-NMDARs were recently shown to be dominantly expressed at posteromedial thalamocortical inputs from layer VI cortical neurons (Topolski et al., 2024). In rodents, the posterior ventrobasal thalamic region is hypothesized to be a critical node for absence seizure generalization (Crunelli et al., 2020). The SWD phenotype observed in *Grin2b* haploinsufficient rats here could hint at potentially dysregulated transmission dynamics at these corticothalamic synapses. A similar observation was previously shown in *Gria4^-/-^*mice, which exhibit SWDs as a result of reduced AMPAR-mediated excitatory postsynaptic currents at cortico-nRT synapses (Paz et al., 2011). Finally, evidence suggests nRT neuronal excitability and communication between the relay thalamus and nRT depend, at least in part, on GluN2B-containing NMDARs (Astori & Lüthi, 2013; Crabtree et al., 2013), the partial absence of which may serve as an alternative or complementary mechanism contributing to seizure generation in *Grin2b^+/-^* animals.

Case studies and parental surveys report a high incidence (>60%) of dysfunctional sleep in individuals with pathological *GRIN2B* variants (Freunscht et al., 2013; Platzer et al., 2017; Simons Searchlight, 2021). Nonetheless, quantitative studies that systematically assess sleep in patients with *GRIN2B* mutations are lacking, thus the evidence of dysfunctional sleep in *Grin2b^+/-^* rats presented here offers valuable insights into the potential sleep impairment phenotypes that may also be found in patient populations. REM, NREM and wake brain states were all significantly altered in *Grin2b^+/-^* rats. Despite a slight increase in the average duration of individual REM epochs, there was a striking reduction in total REM minutes over the 24-hour cycle. The reduced net REM was underpinned by a lower occurrence of REM bouts in *Grin2b* heterozygous knockouts and may indicate altered REM initiation mechanisms. Total NREM minutes were unchanged over 24 hours but *Grin2b^+/-^* rats had fewer NREM bouts that were longer in duration. Lastly, wake total minutes over 24 hours were reduced in mutants, which was mediated by a decrease in the average bout duration despite an increase in the number of bouts. Furthermore, the presence of absence seizures, which primarily occur during wake also reduce the time spent in this state.

NMDAR-dependent signalling plays a critical role in sleep modulation and diurnal rhythmicity. Blocking NMDAR transmission leads to sleep deficits (Stone et al., 1992), while activation of these receptors modulates light-dark locomotor and social behaviour transitions related to sleep-wake physiology (Burgdorf et al., 2019). It was recently shown that phosphorylation of GluN2B serves as a molecular signature of sleep pressure that modulates sleep cycles (Z. Wang et al., 2018). Moreover, REM sleep is particularly impacted by NMDAR processes, and GluN2B antagonism alters the expression of REM sleep by disrupting gamma oscillations (Campbell & Feinberg, 1999; Kocsis, 2012). Given the widespread distribution of GluN2B-NMDARs across various forebrain structures, the observed REM sleep deficits and altered sleep phenotype in *Grin2b^+/-^* rats, likely result from disruptions to molecular sleep signalling and impairment in NMDAR processes across multiple neural circuits.

Recently, NMDARs in the ventrolateral preoptic nucleus of the hypothalamus were shown to be essential for NREM and REM sleep, demonstrating the highest amount of activity during REM (Lu et al., 2000, 2002; Miracca et al., 2022). Pan-neuronal knockout of NMDARs in these cells significantly reduced both NREM and REM sleep, while reduced NMDAR expression in GABAergic populations produced severe NREM fragmentation (Kroeger et al., 2018; Miracca et al., 2022). These observations are consistent with the decreased time in REM sleep in *Grin2b^+/-^* rats. Indeed, it has been suggested that tonic neuronal excitation reliant on extrasynaptic NMDARs (Sah et al., 1989), which are predominantly composed of GluN2B subunits (Tovar & Westbrook, 1999), serves to stabilize hypothalamic sleep-promoting cell firing activity (Miracca et al., 2022). Thus, signalling mediated by GluN2B containing NMDARs could be disrupted in the ventrolateral preoptic nucleus of *Grin2b^+/-^* rats with altered sleep activity. Notably, GluN2B subunits are expressed in hypothalamic regions (L. M. Wang et al., 2008), however, subunit expression patterns specific to the preoptic area have not yet been examined.

Mechanistically, absence seizures are thought to also be associated with insufficiency of sleep promoting mechanisms, as reciprocally interconnected arousal and sleep-promoting neuronal groups control thalamocortical excitability (Danober et al., 1998; Snead, 1995). Reduced activity of sleep promoting GABAergic ventrolateral and median preoptic nuclei were previously shown in epileptic WAG/Rij rats, while inappropriately timed activation of median preoptic nucleus during waking, although insufficient to fully suppress the arousal system, induces SWDs in non-epileptic rats (Suntsova et al., 2009). The observation that absence seizures in *Grin2b*^+/-^ rats are more likely to initiate during wake-NREM transitions, compared to wild-type controls, further supports the idea that preoptic nuclei transmission may be abnormal in these animals. Absence epilepsy is associated with deficits in REM sleep, increased latencies to REM sleep onset, sleep fragmentation and decreased sleep efficiency (Baldy-Moulinier, 1992; Maganti et al., 2005; Saad et al., 2019). Similarly, WAG/Rij rats present with SWDs and have altered sleep patterns consisting of prolonged transitional periods from wake to sleep, lengthened intermediate stages of sleep, more frequently followed by arousals, and decreased time in REM sleep (Coenen & van Luijtelaar, 2003; Gandolfo et al., 1990; van Luijtelaar & Bikbaev, 2007). These findings are in alignment with the sleep and SWD phenotypes observed in *Grin2b^+/-^* rats. Based on the above observations and the findings that NMDARs in the ventrolateral preoptic region are essential for NREM and REM sleep (Lu et al., 2000, 2002; Miracca et al., 2022), it is possible that loss of sufficient GluN2B expression impacts on and dysregulates the circuit activities of basal hypothalamic sleep-promoting structures. Interestingly, the ventrolateral preoptic nucleus receives direct excitatory drive from layer V pyramidal cells, the activity of which is reliant on GluN2B-NMDAR mediated transmission, strengthening the assumption that activity in the preoptic area may also be altered downstream. These scenarios will be interesting to explore in *Grin2b^+/-^* rats, especially considering the reduced REM, altered NREM dynamics, and presence of SWDs in these animals.

Despite the above, we found no correlation between sleep abnormalities and total SWD activity. REM sleep is not critically dependent on thalamocortical circuitry, and while this network is recruited to generate NREM spindles and pace slow oscillatory activity, NREM sleep is also dependent on multiple other hypothalamic and midbrain structures that operate through distinct neurophysiological mechanisms. Given this, the lack of correlation between sleep abnormalities and SWDs may be due to the loss of GluN2B affecting thalamocortical seizure and sleep circuits separately. Such effects on network activity potentially vary depending on the spatial expression of GluN2B.

REM and NREM sleep are dependent on homeostatic sleep drive, with the expression of REM sleep also being modulated by NREM sleep dynamics (Franken & Dijk, 2024). We found no differences in the overall amount of time *Grin2b^+/-^* rats spent in NREM sleep across the 24-hour day, indicating that although NREM sleep dynamics differed between the two genotypes, the homeostatic system responses governing the amount of NREM sleep are likely functionally intact or compensated for in these animals. In contrast, REM sleep was decreased in *Grin2b^+/-^* rats. An interesting study found that the probability of NREM to REM sleep transitions is reduced particularly after long REM sleep periods (>1 min) in rats (Bassi et al., 2009). REM bout duration was statistically longer in *Grin2b^+/-^* rats, with an average length of approximately 65-70 seconds. Therefore, the increased length of REM sleep bouts, which may be attributed to a homeostatic response to reduced REM sleep bouts, could paradoxically also impair the propensity and sleep drive for future REM initiation. The specific factors contributing to reduced REM sleep in *Grin2b^+/-^*animals remain unclear. From a network level perspective, REM sleep is mediated by reciprocally connected nuclei in the pons and brainstem, amongst other regions, which serve to promote NREM-REM transitions, REM sleep muscle atonia, and initiate downstream hippocampal theta rhythms and desynchronizes cortical activity (Boissard et al., 2002; Park & Weber, 2020; Scammell et al., 2017). It is plausible that in *Grin2b^+/-^* mutant animals, signalling irregularities are present within REM promoting circuits, particularly those relating to NREM suppression and REM sleep initiation. Alternatively, downstream seizure related processes, such as imbalance of neuromodulators or disruption of regular circadian patterns may also account for reduced REM sleep in *Grin2b^+/-^* rats (Bazil et al., 2000).

Our study demonstrates altered spectral power in *Grin2b^+/-^* animals during SWDs and NREM sleep. In *Grin2b^+/-^*rats the spectral slope of SWD epochs showed a peak frequency in the theta range with strong resonant harmonics in higher frequency bands. It is possible this harmonic is related to the prolonged SWD durations observed in *Grin2b^+/-^* rats, as longer seizures may allow for greater accumulation of synchronized neuronal activity. Delta power during seizures in *Grin2b^+/-^* animals was also significantly higher, and previous work in GAERS and WAG/Rij rats has described increases in delta preceding SWD onset as pro-epileptogenic (Bosnyakova et al., 2007; van Luijtelaar et al., 2011). SWDs predominantly occur during low-vigilance states such as drowsiness and light slow-wave sleep, which are characterized by EEG slowing. Mechanistically, the pro-epileptic nature of an increased delta amplitude may relate to burst firing in thalamocortical and corticothalamic neurons, a pattern associated not only with SWDs but also with sleep spindles and NREM delta and slow wave activity, which is governed by the degree of hyperpolarization of these cells (Crunelli & Hughes, 2010; McCormick & Bal, 1997; Steriade et al., 1993). During NREM sleep *Grin2b^+/-^* animals had a pronounced increase in beta power, which is seen as a reliable marker of cortical hyperarousal and low quality sleep in individuals with sleep disorders and insomnia (Fernandez-Mendoza et al., 2016; Shi et al., 2022; Spiegelhalder et al., 2012). Beta oscillations are believed to arise from intra-cortical connectivity concurrent with strong distal inputs to cortex, possibly of thalamic origin, and are associated with the synchronization of neural activity across different brain regions (Benedek et al., 2016; Sherman et al., 2016). NMDAR antagonists increase beta power (Rebollo et al., 2018; van Lier et al., 2004). Similarly, NMDAR hypofunction in schizophrenia, associated with *GRIN2B* variants, elevates resting-state beta (Rebollo et al., 2018; Takasaki et al., 2016; Venables et al., 2009), indicating beta rhythmicity depends on intact NMDAR function. Our data could therefore indicate potentially compromised cortical signalling, altered connectivity and dysregulated synchronization properties in *Grin2b^+/-^*animals, particularly in relation to corticothalamic signalling, which is recruited during both SWDs and NREM sleep.

Finally, we found that SWDs in *Grin2b^+/-^*rats are sensitive to systemic treatment with ethosuximide and memantine. Ethosuximide is one of most widely used drugs for the treatment of absence seizures, and as in this study, successfully diminishes SWDs in rodent models and patients (Glauser et al., 2013; Penovich & James Willmore, 2009; Richards et al., 2003; Taylor et al., 2019), mainly through its antagonistic effect on T-type calcium channels. In contrast, memantine had no effect on the frequency of occurrence of SWDs, but did however, decrease the average duration of seizures. This effect may be due to memantine’s action at the NMDAR magnesium binding site, which is only exposed during periods of membrane depolarization (H. Chen et al., 1992; H. S. Chen & Lipton, 1997). Consequently, absence seizure initiation may remain unaffected but, after successive bouts of membrane depolarization during a SWD, the events may shorten when memantine blocks the receptor at the magnesium site. Extending these findings, we also show that direct infusion of ethosuximide or memantine into the nRT acutely reduced SWD occurrence, likely by shifting thalamic firing output from burst to tonic mode via blockade of T-type calcium channels and NMDARs. This demonstrates that, as in other rodent models (Cao et al., 2020; Koerner et al., 1996; McCafferty et al., 2018; Richards et al., 2003), the thalamus is critically engaged in SWD generalization and maintenance in *Grin2b^+/-^* rats. The effectiveness of memantine on drug-resistant early-onset epilepsy in individuals with gain of function *GRIN2* variants (*GRIN2A, GRIN2B* and *GRIN2D*) is mixed (Li et al., 2016; Pierson et al., 2014; Platzer et al., 2017; XiangWei et al., 2018) and previously, memantine treatment did not improve seizure frequency in four patients with *GRIN2B* gain of function mutations.(Platzer et al., 2017). Moreover, memantine is rarely used in patients with loss of function mutations, although this is consistent with the current understanding of NMDAR pharmacology and the differential effects of gain versus loss of function mutations on glutamatergic signalling. With the above in mind, our data suggest that memantine could have favourable outcomes with respect to seizure manifestations in *Grin2b* haploinsufficiency. However, the potential effects of memantine on seizures associated with *GRIN* mutations resulting in NMDAR hypofunction, or its use as an adjunctive therapy require further research and clinical evaluation before any firm conclusions can be drawn.

In conclusion, we report on the presence of absence seizures and abnormal sleep architecture in a novel rat model of heterozygous *Grin2b* knockout. The model offers the first reliable spontaneous seizure phenotype and pre-clinical quantitative evidence of dysfunctional sleep relating to *GRIN2B* haploinsufficiency. Additionally, we demonstrate that seizures in *Grin2b^+/-^*animals are sensitive to ethosuximide and memantine, and propose both drugs could favourably impact on absence seizure severity. It is important to note, however, that absence seizures have not been reported in loss-of-function patients, which should be considered when assessing the clinical relevance of the model. Overall, the data presented here are consistent with clinical observations and as such provide a translationally relevant tool for future research that should aim at defining the underlying pathological mechanisms and developing targeted therapeutics for *GRIN2B* related disorders.

## Data availability

Raw data are available upon request from the corresponding author. The source code files for the offline sleep state classification and SWD detection algorithms are accessible at https://github.com/Gonzalez-Sulser-Team/AUTOMATIC-SLEEP-SCORER and https://github.com/Gonzalez-Sulser-Team/SWD-Automatic-Identification, https://zenodo.org/records/12700972.

## Abbreviations

ASD: autism spectrum disorder
NMDAR: N-methyl-D-aspartate receptor
SWD: spike and wave discharge
REM: rapid eye movement sleep
NREM: non-rapid eye movement sleep
nRT: reticular thalamic nucleus.

