## Supplementary Data for "Absence seizures and sleep-wake abnormalities in a rat model of *GRIN2B* neurodevelopmental disorder"

### Content

**Supplementary Table 1**

**Supplementary Table 2**

**Supplementary Figure 1**

**Supplementary Figure 2**

**Supplementary Figure 3**

**Supplementary Figure 4**

**Supplementary Figure 5**

**Supplementary Figure 6**

**Supplementary Figure 7**

| | ***Visually scored epochs*** | | |
|---|---|---|---|
| | *REM* | *NREM* | *Wake* |
| **Automatically Scored Epochs** | | | |
| *REM* | **2378** | 107 | 233 |
| *NREM* | 119 | **20637** | 781 |
| *Wake* | 201 | 285 | **19254** |
| *Total* | 2698 | 23594 | 20268 |
| *Agreement (%)* | 88.1 | 87.5 | 95 |
| *Global agreement (%)* | 90.8 | | |
| *Cohen's kappa (κ) (± $Se_κ$)* | 0.83 ± 0.002 | | |

**Supplementary Table 1: Comparison of the performance of sleep-wake automated scoring algorithm and visual scoring with results from statistical comparison.** Numbers represent epochs scored visually or automatically for REM, NREM and wake states for all animals in sleep-wake analysis. The diagonal values (bold and grey background) represent instances of agreement between both methods, meaning both produced the same scored state output. None-diagonal numbers represent instances of disagreement in state scoring. Total epochs scored, agreement percentage, global agreement across states and Cohen’s kappa values are shown below scored epoch values.

| | *Effect of genotype* | | | *Effect of sex* | | | *Effect of genotype x sex* | | |
|---|---|---|---|---|---|---|---|---|---|
| | *F* | *DF* | *P* | *F* | *DF* | *P* | *F* | *DF* | *P* |
| ***Total REM duration*** | 8.23 | 1 | 0.0095 | 0.0094 | 1 | 0.92 | 0.83 | 1 | 0.37 |
| ***Number of REM bouts*** | 12.46 | 1 | 0.0021 | 0.033 | 1 | 0.86 | 0.39 | 1 | 0.54 |
| ***Average REM duration*** | 2.49 | 1 | 0.13 | 1.76 | 1 | 0.41 | 0.036 | 1 | 0.85 |
| ***Total NREM duration*** | 0.0074 | 1 | 0.93 | 0.46 | 1 | 0.5 | 0.064 | 1 | 0.8 |
| ***Number of NREM bouts*** | 2.96 | 1 | 0.101 | 0.0066 | 1 | 0.94 | 0.0045 | 1 | 0.95 |
| ***Average NREM duration*** | 0.28 | 1 | 0.11 | 0.83 | 1 | 0.37 | 0.0025 | 1 | 0.96 |
| ***Total wake duration*** | 5.054 | 1 | 0.036 | 1.66 | 1 | 0.21 | 0.12 | 1 | 0.73 |
| ***Number of wake bouts*** | 7.13 | 1 | 0.015 | 3.77 | 1 | 0.066 | 0.025 | 1 | 0.62 |
| ***Average wake duration*** | 10.38 | 1 | 0.0043 | 5.69 | 1 | 0.027 | 1.9 | 1 | 0.18 |
| | | | | | | | ***Effect of genotype x sex x hour*** | | |
| | | | | | | | *F* | *DF* | *P* |
| ***REM hour-by-hour*** | 8.84 | 1 | 0.0075 | 0.85 | 1 | 0.037 | 1.24 | 1 | 0.21 |
| ***NREM hour-by-hour*** | 0.043 | 1 | 0.52 | 0.45 | 1 | 0.5 | 1.46 | 1 | 0.078 |
| ***Wake hour-by-hour*** | 1.46 | 1 | 0.038 | 5.054 | 1 | 0.036 | 1.31 | 1 | 0.16 |

**Supplementary Table 2: Values for statistical analysis across brain states, genotypes and sex.** Left most column contains description of test statistic and measurements analysed. Columns 2–4 contain relevant *F, DF* and *P* values for the effects of genotype and sex, and interaction effect between them. Values in rows 3–11 correspond to results from Two-way ANOVA test statistics and values in rows 14–16 correspond to results from Linear mixed models used in hour-by-hour time course analysis of sleep-wake distribution. Statistical results relate to Supplementary Fig. 7.

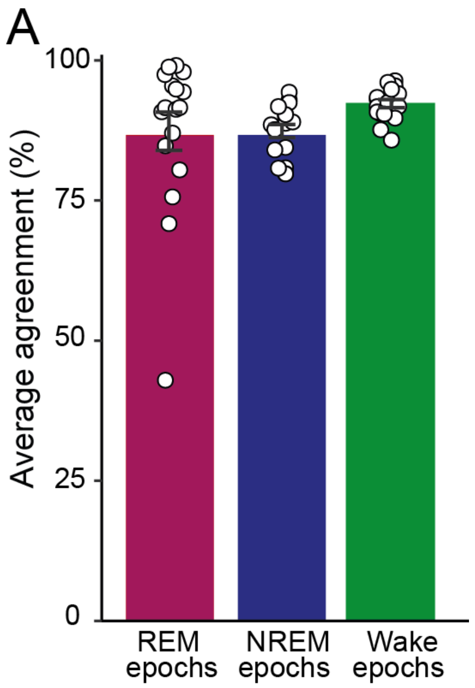


**Supplementary Figure 1: Percentage Agreement Between Sleep-Wake Automated Scoring Algorithm and Visual Scoring.** Agreement between visually and automatically scored epochs of REM, NREM and wake states is 88.1%, 87.5% and 95% respectively. Overall agreement between the scoring methods is 90.8%. The kappa coefficient was 0.83 (± 0.002 SE). See also Supplementary Table 1 for further details. Bars indicate mean values (mean ± SEM). Points correspond to values from individual rats.

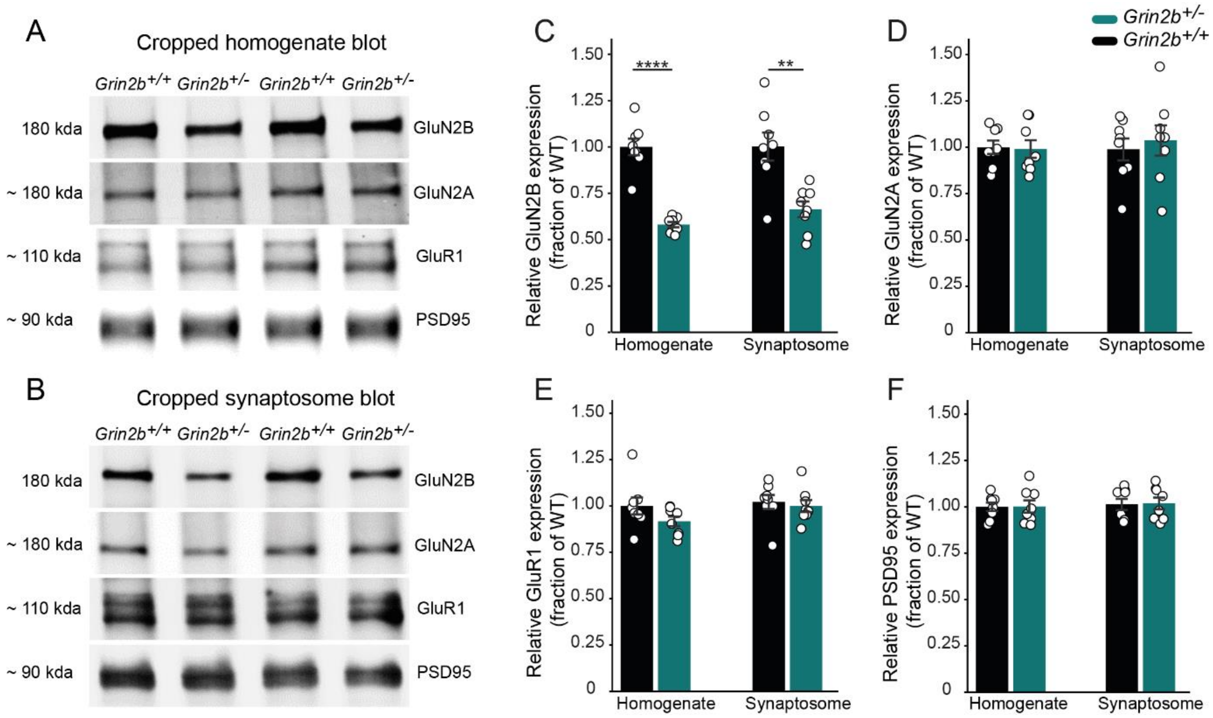


**Supplementary Figure 2: *Grin2b* deletion in rats results in reduction of endogenous GluN2B expression in hippocampus.** Representative Western blots of extracts from rat hippocampal brain (**A**) homogenates and (**B**) synaptosomes, full length blots in Supplementary Fig 3. Bands in the molecular weight range expected for full length GluN2B, GluN2A, GluR1 and PSD95 were detected in homogenates and synaptosomes from wild-type and *Grin2b*$^{+/-}$ animals. (**C**) Quantification of GluN2B protein from homogenates and synaptosomes reveals a significant decrease in *Grin2b*$^{+/-}$ rats (homogenate $P$ < 0.00001, synaptosome $P$ 0.0016, Two-sample unpaired t-tests). There was no change in *Grin2b*$^{+/-}$ rats in expression levels of (**D**) GluN2A ($P$ = 0.88, $P$ = 0.64, Two-sample unpaired t-test), (**E**) GluR1 ($P$ = 0.14, $P$ = 0.67, Two-sample unpaired t-tests), and (**F**) PSD95 ($P$ = 0.96, $P$ = 0.71, Two-sample unpaired t-test and Wilcoxon rank sum tests). Detailed statistics in text. Bars indicate mean values (mean ± SEM). Points correspond to values from individual rats.

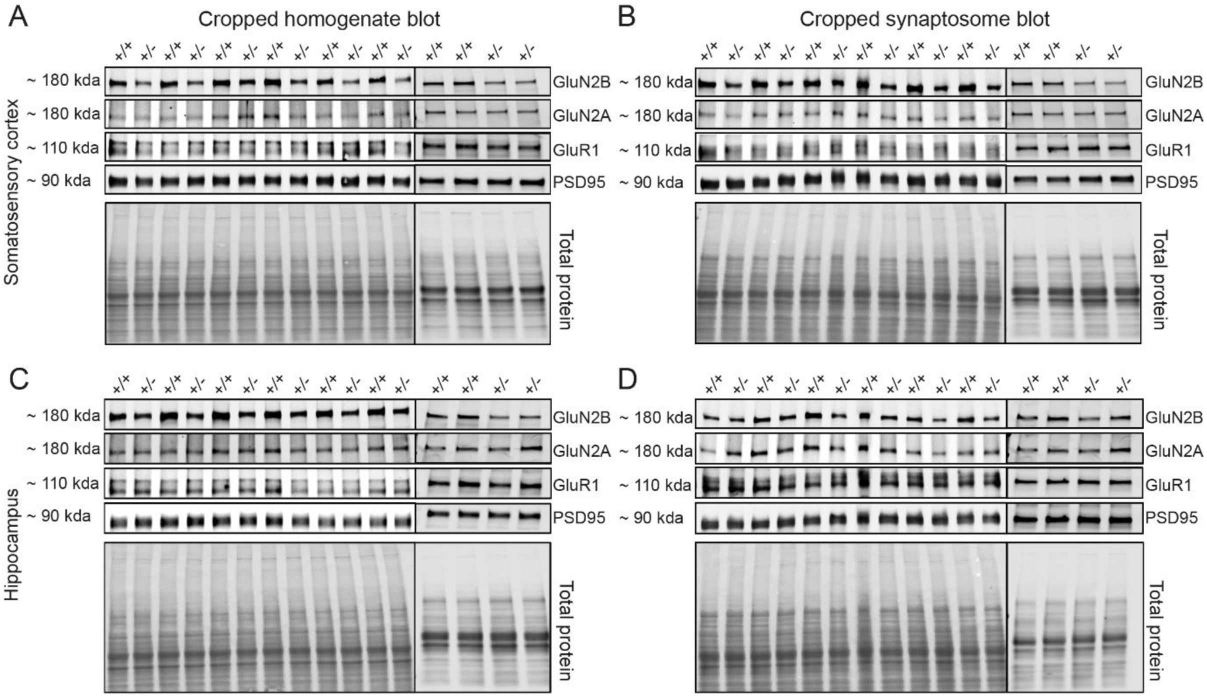


**Supplementary Figure 3: *Grin2b* deletion in rats results in reduction of endogenous GluN2B expression in somatosensory cortex and hippocampus.** Entire set of Western blots of extracts from rat somatosensory brain (**A**) homogenates and (**B**) synaptosomes, and extracts from rat hippocampal brain (**C**) homogenates and (**D**) synaptosomes. Bands in the molecular weight range expected for full length GluN2B, GluN2A, GluR1 and PSD95 were detected in homogenates and synaptosomes from wild-type and *Grin2b*$^{+/-}$ animals.

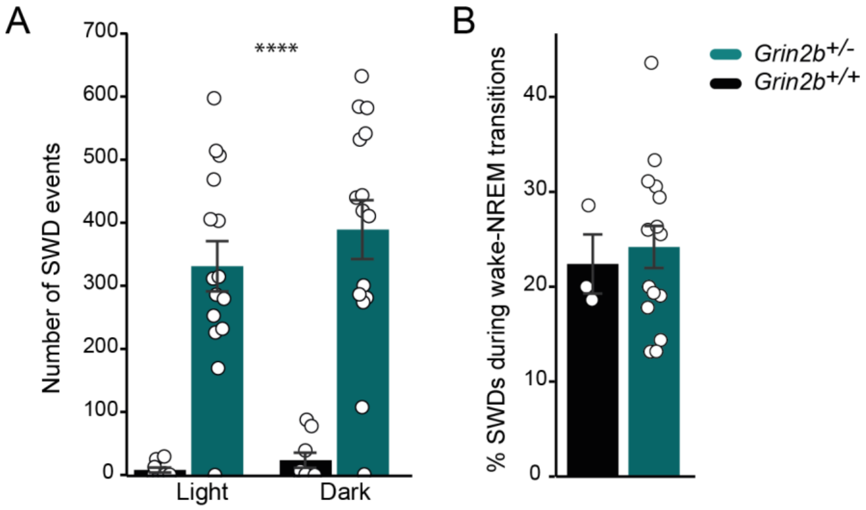


**Supplementary Figure 4: Light phase did not influence the prevalence of SWDs and the percentage of wake-NREM transitions did not differ between genotypes.** (**A**) Number of SWD events during light and dark phases did not significantly differ between light and dark phases but were significantly increased in *Grin2b*$^{+/-}$ rats (effect of genotype P < 0.00001; effect of phase P = 0.28; phase x genotype P = 0.53, Linear Mixed Model), (*** = effect of genotype). *Grin2b*$^{+/-}$ rats showed similar SWD event numbers in both light and dark periods, which significantly exceeded SWD amounts in wild-type littermates during both phases (effect of genotype $P < 0.00001$; effect of phase $P = 0.28$; phase x genotype $P = 0.53$, Linear Mixed Model). Bars indicate mean values (mean ± SEM). Points correspond to values from individual rats. (**B**) A higher proportion of SWDs in NREM initiate during wake-NREM transitions in *Grin2b*$^{+/-}$ animals when compared to wild-type animals ($P = 0.74$, Two-sample unpaired t-test). Bars indicate mean values (mean ± SEM) and points correspond to values from individual rats. Amount of SWDs initiating at wake-NREM transitions were not different between *Grin2b*$^{+/-}$ rats and *Grin2b*$^{+/+}$ wild-types that did have SWDs at transitional periods (3/9 animals) ($P$ = 0.74, Two-sample unpaired t-test).

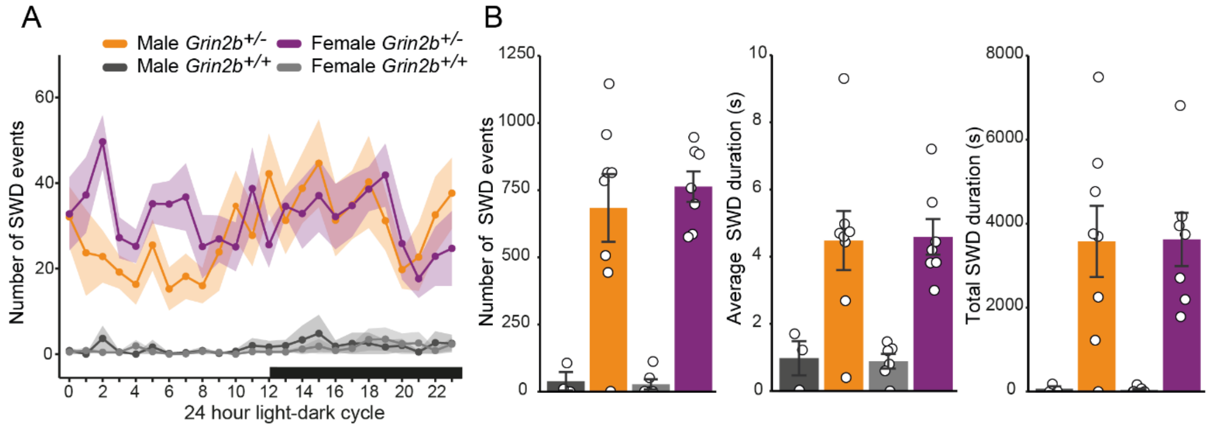


**Supplementary Figure 5: No differences in SWD properties between male and female *Grin2b$^{+/-}$* rats.** (**A**) Number of SWD events plotted by hour over the 24-hour light-dark cycle with black bar on x-axis indicating lights off in the animal facility. Hour of the day and sex do not impact on the amount of SWDs in *Grin2b$^{+/-}$* male and female rats, which consistently had significantly more seizures than wild-type sex-matched littermates (hour $P = 0.021$, genotype $P = 0.00001$, sex $P = 0.73$, hour x genotype x sex $P = 0.66$, Linear Mixed Model). Points indicate mean values of all animals (mean ± SEM). (**B**) Bar plots of total number of SWDs events (left), SWD average duration (middle) and total SWD duration (right). *Grin2b$^{+/-}$* rats (SWD events sex $P = 0.74$, genotype $P < 0.0001$, sex x genotype $P = 0.66$; average duration sex $P = 0.99$, genotype $P = 0.0001$, sex x genotype $P = 0.89$; total duration sex $P = 0.99$, genotype $P = 0.0001$, sex x genotype $P = 0.96$, Two-way ANOVA tests). Bars indicate mean values (mean ± SEM). Note: we do not indicate significance (*) where effect from genotype is found. Points correspond to values from individual rats. Detailed statistic can be found in results text.

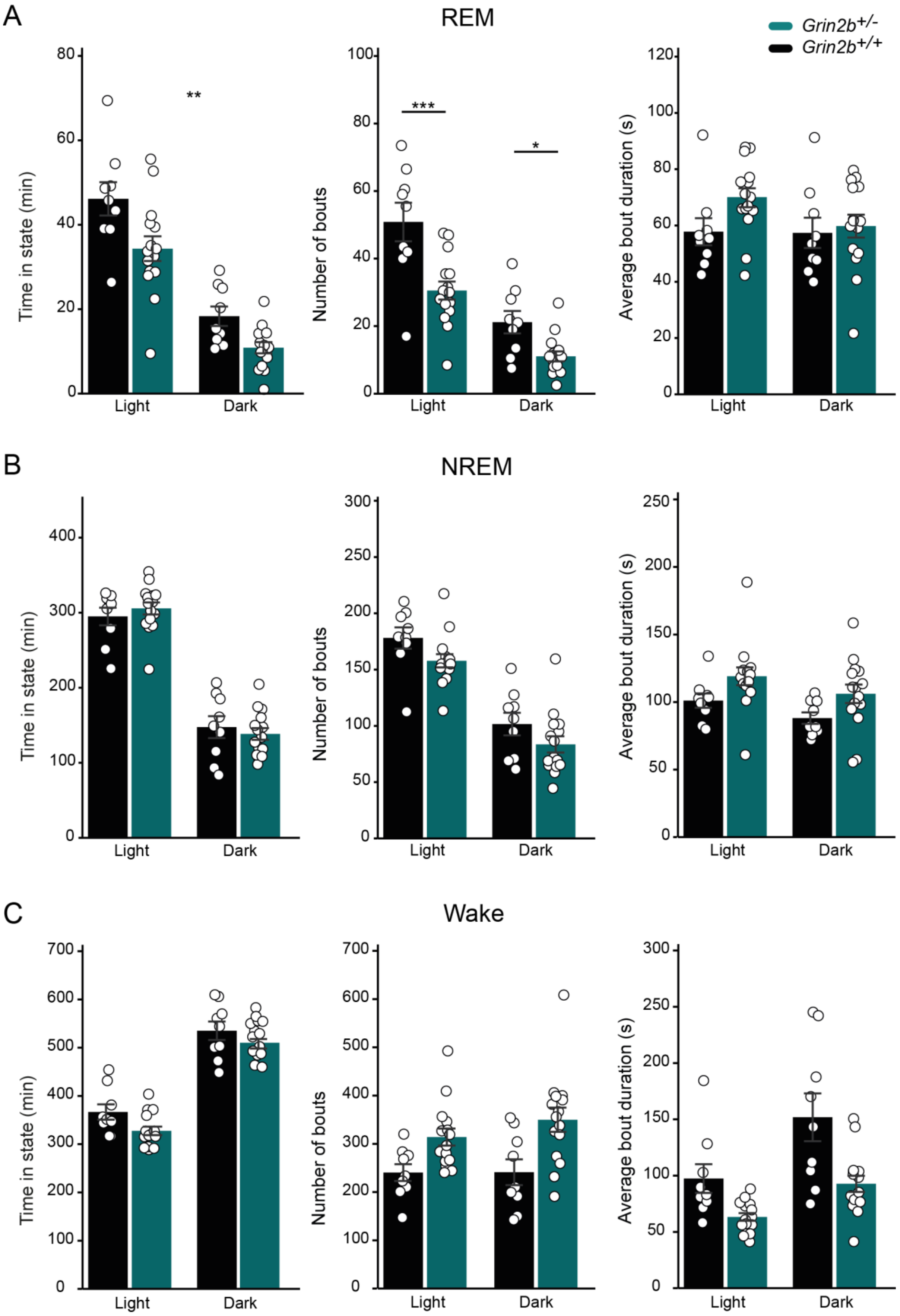


**Supplementary Figure 6: Reduced REM sleep in *Grin2b*$^{+/-}$ rats during the light and dark periods.** Quantification showing total time (left), number of bouts (middle) and average bout duration (right) for (**A**) REM, (**B**) NREM and (**C**) wake during the 12-hour light and dark periods. Bars indicate mean values (mean ± SEM) and points correspond to values from individual rats. Total time in REM sleep was overall reduced in *Grin2b*$^{+/-}$ rats compared to wild-types, and reduced REM sleep was not specific to either light or dark phases (phase $P$ < 0.00001, genotype $P$ = 0.0046, phase x genotype $P$ = 0.37, Linear Mixed Model). In comparison to *Grin2b*$^{+/+}$ animals, the number of REM bouts in *Grin2b*$^{+/-}$ rats was reduced, both during the light and the dark periods, while no differences were found between the two groups in the average length of REM bouts (REM bouts phase $P$ < 0.00001, genotype $P$ = 0.00067, phase x genotype $P$ = 0.045 (light $P$ = 0.0001, dark $P$ = 0.032); REM bout duration REM phase $P$ = 0.13, genotype $P$ = 0.18, phase x genotype $P$ = 0.16; Linear Mixed Models with Tukey *post hoc* tests). Analysis by light-dark phase revealed no difference between the two genotypes for total time spent in NREM sleep, the number of NREM bouts and the average NREM bout durations (Time in NREM phase $P$ < 0.00001, genotype $P$ = 0.94, phase x genotype $P$ = 0.34; NREM bouts phase $P$ < 0.00001, genotype $P$ = 0.07, phase x genotype $P$ = 0.84; NREM bout duration phase $P$ = 0.00055, genotype $P$ = 0.055, phase x genotype $P$ = 0.99; Linear Mixed Models). Relative to wild-type littermates, *Grin2b*$^{+/-}$ rats had fewer total minutes of wake, and more wake bouts of increased duration, however, differences were irrespective of the light or dark periods (Time in wake phase $P$ < 0.00001, genotype $P$ = 0.013, phase x genotype $P$ = 0.64; wake bouts phase $P$ = 0.25, genotype $P$ = 0.0049, phase x genotype $P$ = 0.27; wake bout duration phase $P$ < 0.00001, genotype $P$ = 0.0015, phase x genotype $P$ = 0.14; Linear Mixed Models). Detailed statistic can be found in results text.

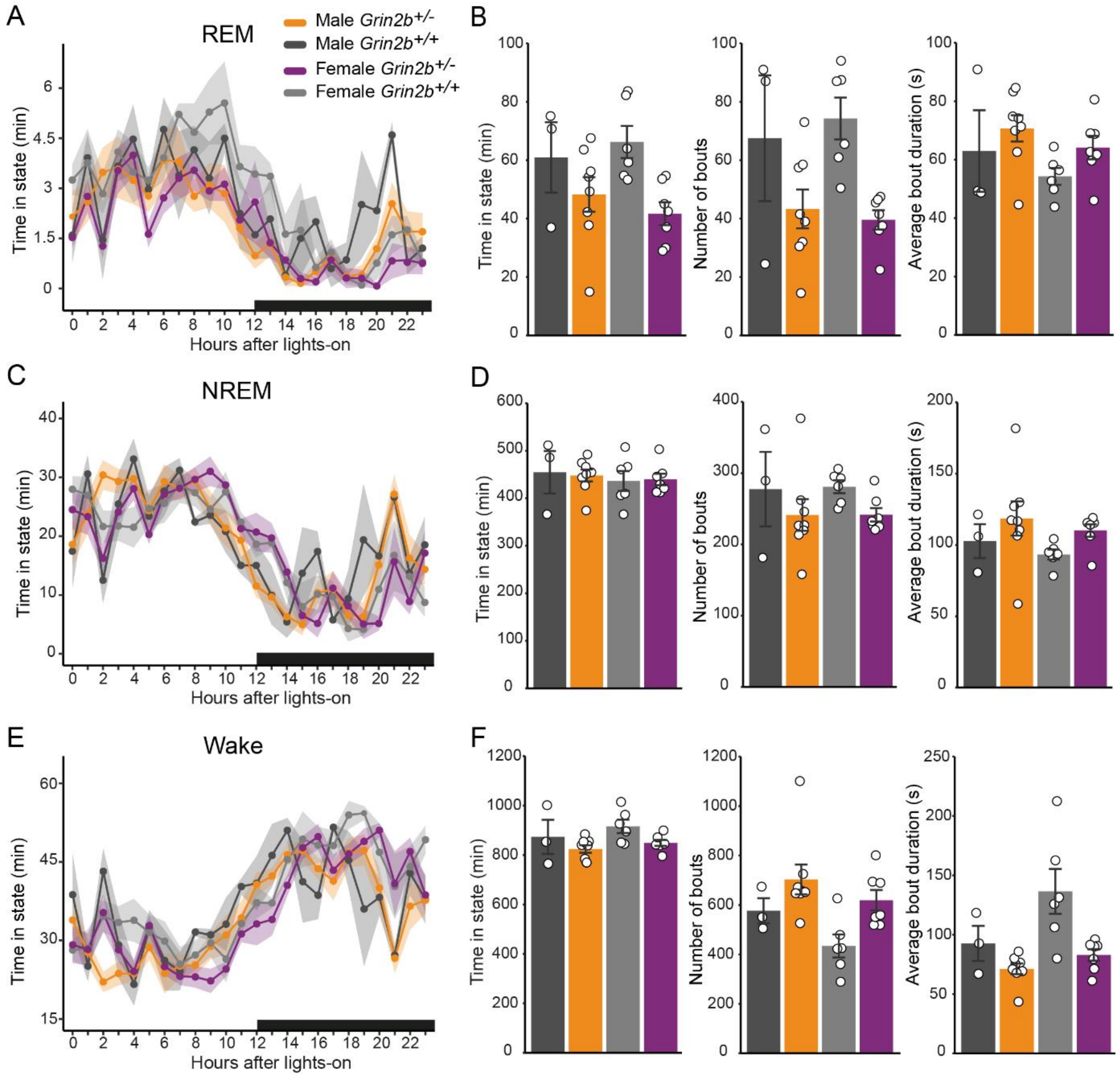


**Supplementary Figure 7: Sleep-wake physiology in *Grin2b$^{+/-}$* animals does not differ between male and female animals.** Quantification showing the time spent in (**A**) REM, (**C**) NREM and (**E**) wake by hour across the 24-hour light-dark cycle; points indicate mean values of all animals (mean ± SEM). Bar plots showing total time (left), number of bouts (middle) and average bout duration (right) for (**B**) REM, (**D**) NREM and (**F**) wake during the full 24 hours; bars indicate mean values (mean ± SEM) and points correspond to values from individual rats. (**A**) Time in REM is reduced in both female and male *Grin2b$^{+/-}$* rats relative to female and male wild-type littermates, however, this difference is not due to animals' sex or to specific hours of the day. (**B**) For the full 24 hours, sex did not impact on total REM sleep or REM bouts, as each was reduced in both male and female *Grin2b$^{+/-}$* mutants relative to their wild-type sex-matched littermates. Average REM bout duration was not different between both sexes and genotypes. (**C**) Time in NREM is similar between *Grin2b$^{+/-}$* and wild-type animals of both sexes throughout each hour of the day. (**D**) For the 24-hour day, sex did not impact on total NREM sleep, NREM bouts and average NREM bout duration, and these metrics did not differ between male and female *Grin2b$^{+/-}$* and *Grin2b$^{+/+}$* animals. (**E**) Time in wake is reduced in both female and male *Grin2b$^{+/-}$* rats relative to female and male *Grin2b$^{+/+}$* littermates, and this difference is not due to animals' sex or to specific hours of the day. (**F**) There we no sex dependent differences for the total wake time, number of wake bouts and average wake bout duration, when quantified for the full 24 hours. For detailed statistics see Supplementary Table 2. Note: we do not indicate significance (*) where effect from genotype is found. Bars indicate mean values (mean ± SEM). Points correspond to values from individual rats.
